# A copper-dependent, redox-based hydrogen peroxide perception in plants

**DOI:** 10.1101/2025.07.22.666036

**Authors:** Nobuaki Ishihama, Yohta Fukuda, Yumiko Shirano, Kaori Takizawa, Ryoko Hiroyama, Kazuhiro J. Fujimoto, Hiroki Ito, Mayumi Nishimura, Takeshi Yanai, Tsuyoshi Inoue, Ken Shirasu, Anuphon Laohavisit

## Abstract

Redox-related molecules, such as quinones and reactive oxygen species (ROS), are important signaling molecules for all living organisms. A plant-specific leucine rich-repeat receptor-like kinase (LRR-RLK) CANNOT RESPOND TO DMBQ 1 (CARD1), also known as HYDROGEN-PEROXIDE-INDUCED Ca2+ INCREASES (HPCA1), perceives both quinones and ROS, but knowledge on key structural features is unknown. Here, we determine the structure of the CARD1 ectodomain and uncover its unique features. Structural studies, coupled with genetics and biochemical analysis, demonstrated that previously identified unique cysteine residues are not essential for signal perception in CARD1. Interestingly, CARD1 harbors a copper ion on the surface of the ectodomain *via* histidine-coordination that is crucial for hydrogen peroxide signaling. Our work reports the first copper-dependent redox receptor and highlights a unique perspective between a receptor and non-peptide stimuli during their perception.

## Main

Plants’ ability to perceive external and internal signals via an extensive number of cell surface receptors reinforce their adaptation to fluctuating environment. Among these receptors, the leucine-rich repeat receptor-like kinases (LRR-RLKs) represents one of the largest and most rapidly expanding gene families in the plant kingdom^1^. These LRR-RLKs are highly diversified and are responsible for perceiving a wide range of signals, from developmental cues to pathogen recognition^2^. While many plant LRR-RLKs perceive their cognate proteinaceous ligands, a notable exception is the brassinosteroid receptor BRI1, which perceives a steroid ligand^3^.

The *CANNOT RESPOND TO DMBQ 1* (*CARD1*) receptor (also known as *HYDROGEN-PEROXIDE-INDUCED Ca2+ INCREASES* – *HPCA1*) was previously identified to perceive quinone compounds^4^, particularly the 2,6-dimethoxy-1,4 benzoquinone (DMBQ), as well as reactive oxygen species (ROS) such as hydrogen peroxide (H_2_O_2_)^5^. CARD1 features a unique cluster of cysteine residues (Cys395, Cys405, Cys421, Cys424, Cys434, and Cys436) at the C-terminal part of its ectodomain, raising the possibility that these residues may be involved in the perception of both quinones and ROS^4,5^. Previous studies demonstrated that mutation in cysteine residues Cys395 and Cys405 resulted in a loss of Ca^2+^ signaling in response to DMBQ. However, it remains unclear whether such effect is due to the structural role of cysteine residues or their direct involvement in quinone perception^4^. In contrast, cysteine residues Cys421, Cys424, Cys434, and Cys436 have been implicated in H_2_O_2_ sensing, with disulfide bonds formation between Cys421-Cys424 and Cys434-Cys436 proposed as crucial for H_2_O_2_ perception ^5^. Notably, these residues do not appear to affect quinone sensing^4^.

To elucidate the distinct molecular mechanisms underlying quinone and H_2_O_2_ perception by CARD1, we determined the cryo-electron microscopy (cryo-EM) structure of CARD1 ectodomain (EctoCARD1) at 3.26 Å resolution. Our structural analysis revealed previously unrecognized features of EctoCARD1 and, contrary to previous analysis, demonstrating that cysteines residues are not mechanistically required for the perception of either quinone or H_2_O_2_. Instead, we identified a distinct copper binding histidine cluster in the EctoCARD1, that is necessary for H_2_O_2_-induced signaling, but is not essential for quinone perception. Overall, this study delineates how CARD1 distinguishes between quinone and H_2_O_2_ signal perception, and offers the first structural insights into the LRR-RLK clade VIII-1 subfamily. To our knowledge, this is the first evidence of metal ion coordination – specifically copper (Cu) –in the LRR region of a plant receptor. Rather than relying on a typical cysteine-mediated, redox-based perception, CARD1 employs a unique copper-dependent, redox-based for H_2_O_2_ perception in plants.

### H_2_O_2_ perception by CARD1 and its phylogeny

We first validated previous finding that *CARD1* is required for H_2_O_2_-induced Ca^2+^ elevation, using mutants in *Arabidopsis* that expresses aequorin – a Ca^2+^ indicator based on bioluminescence (hereafter, WT-AEQ)^4^. Indeed, all *card1* mutant alleles displayed abolished Ca^2+^ signaling in response to H_2_O_2_ (Supplemental Fig. 1a) while complementation assay restored the phenotype for low H_2_O_2_ (100 µM) and high H_2_O_2_ (5 mM) (Fig. 1a, b), confirming its functional requirement. Kinetic analysis revealed that H_2_O_2_ exhibited a higher apparent *K*_m_ (58.6 µM) compared to DMBQ (1.68 µM^4^), highlighting different sensitivity profiles for ROS and quinones (Supplemental Fig. 1b). Next, we tested if ROS generated in the apoplast, such as by the NADPH oxidases (RBOHD and RBOHF), could be perceived by CARD1. Flg22 peptide is a well-known pathogen-associated molecular patterns (PAMP) molecule that elicits PAMP-triggered immunity that includes NADPH oxidase-mediated ROS production. WT-AEQ seedlings were challenged with 1 µM flg22 peptide and showed two distinct Ca^2+^ peaks (Fig. 1c) as previously reported^6^. Interestingly, in all *card1* alleles, the second Ca^2+^ peak was absent (Supplemental Fig. 1c). If CARD1 was to sense the ROS generated from RBOHD, flg22-induce Ca^2+^ phenotype in *card1* should phenocopies *rbohd*. Indeed, the second peak was abolished in both *card1* and *rbohd* mutants, implying that ROS which is generated by RBOHD during flg22 signaling is perceived by CARD1 in the apoplast (Fig. 1d), further highlighting that CARD1, in addition to quinones, is capable of perceiving ROS generated during plant immunity signaling.

**Figure 1.**
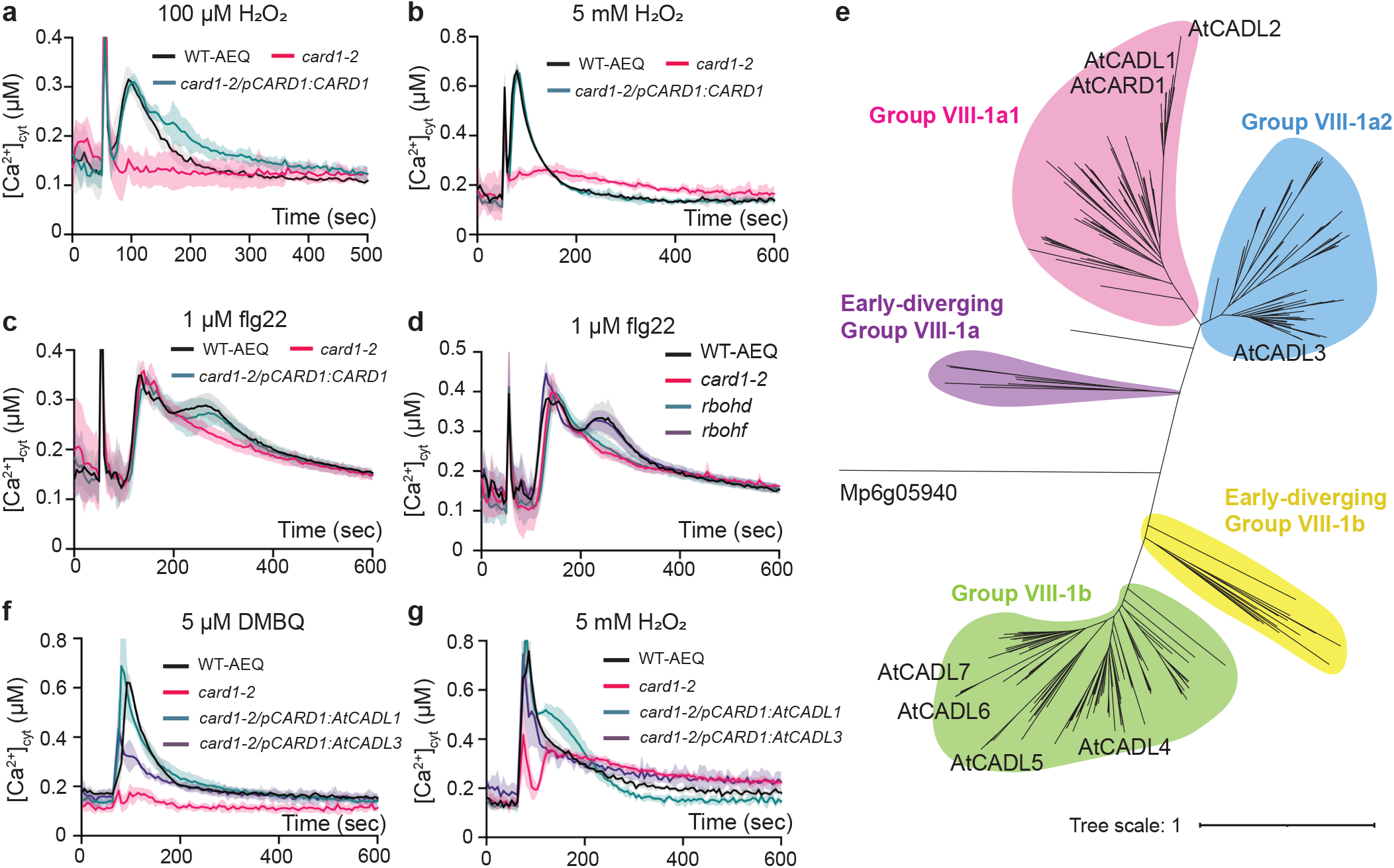
CARD1 perceives apoplastic ROS and CARD1 phylogeny. **a-c**, [Ca^2+^]_cyt_ response of *Arabidopsis* seedlings WT-AEQ, *card1-2* and complemented line (*card1-2/pCARD1:CARD1*) in response to (**a**) 100 µM H_2_O_2_, (**b**) 5 mM H_2_O_2_, and (**c**) 1 µM flg22. Stimuli were applied at 60 s. Data are mean ± SE (*n* = 4) for H_2_O_2_ treatments and (*n* = 6) for flg22. **d**, [Ca^2+^]_cyt_ response of *Arabidopsis* seedlings WT-AEQ, *card1-2, rbohd*, and *rbohf* in response to 1 µM flg22. Stimuli were applied at 60 s. Data are mean ± SE (*n* = 6). *rbohf* was included as a negative control. **e**, An unrooted phylogenetic tree of CARD1 protein in land plants. Amino acid sequences were mined from the Phytozome database version 12.1 and full protein sequences were aligned using MUSCLE (default parameters). See methods for full descriptions.

We next clarify the phylogeny of CARD1 in land plants and we surveyed for CARD1 and CARD1-like (CADLs) homologs across land plants and the late divergent charophyte groups. Following manual curation, 405 CARD1/CADLs sequences, across 57 species, were identified (Supplementary Data 1). These homologs are conserved throughout embryophytes, but are absent in Zygnematophyceae, the sister clade and the closet relative of the embryophytes whose genome has recently been published^7^. Phylogenetic analysis revealed two major subfamilies within Group VIII-1 of LRR-RLK family (VIII-1a and VIII-1b) which diverged early in embryophytes evolution (Fig. 1e). We further identified two distinct subgroups within LRR-RLK group VIII-1a (VIII-1a1 and VIII-1a2). The split of the LRR-RLK VIII-1a to a1 and a2 subgroups did not occur until after the bryophytes (Fig. 1E). CARD1, along with CADL1 and CADL2, belongs to Group VIII-1a1; CADL3 is in Group VIII-1a2; while CADL4 through CADL7 are part of Group LRR-RLK VIII-1b (Fig. 1e). To establish whether the perception of quinones and ROS are observed in other *Arabidopsis* CARD1 homologs in Group VIII-1a, we performed complementation assay by expressing other *Arabidopsis* CARD1 homologs (AtCADL1 and AtCADL3) in *card1-2* plants under CARD1 promoter. Both homologs can restore DMBQ and H_2_O_2_-induced Ca^2+^ signaling (Fig. 1f, g). These findings support the idea that quinone and H_2_O_2_ perception capabilities in CARD1 and CADL homologs are likely not unique to CARD1 alone, and that DMBQ and H_2_O_2_ perception could be performed by other homologs, depending on when and where each gene is expressed.

### Overall EctoCARD1 protein structure

To gain further insights into how CARD1 perceives quinones and ROS, such as DMBQ or H_2_O_2_, respectively, we sought to determine the structure of CARD1 protein by employing previously optimized expression and purification strategies in *Nicotiana benthamiana*^8^. We also fused EctoCARD1 to maltose-binding protein (MBP) to increase the size of the EctoCARD1 to a respectable size and purified the protein (Supplemental Fig. 2) for cryo-EM analysis (Supplemental Fig. 3). The MBP region did not appear in the 2D classification particles, allowing us to reconstruct a cryo-EM map corresponding solely to the EctoCARD1 region (Supplemental Fig. 3). We determined the structure of EctoCARD1 (PDB 9VBD) at 3.26 Å resolution (Fig. 2a, b, Supplementary Table 1). EctoCARD1 has two distinct regions: the N-terminal LRR region and the C-terminal domain, which is connected by a loop region containing conserved Cys residues (Cys421, Cys424, Cys434, and Cys436) (Fig. 2c). The fold of the C-terminal domain is highly similar to the SEA (Sperm protein, Enterokinase, and Agrin domain) (Supplementary Table 2, 3)^9^. The LRR and SEA-like domains interact extensively through hydrogen bonds and electrostatic interactions (Fig. 2d). Notably, the interaction surface is reminiscent of the ligand-binding site in other known LRR-RLK such as FLS2 or HAESA receptors^3^, suggesting that the SEA-like domain may play a role in modulating receptor activity. The LRR domain consists of 14 LRRs, adopting a characteristic twisted solenoid configuration resembling a horseshoe structure^3^. This region also includes a highly conserved cysteine-pair capping domain at both the N terminus (Cys53 and Cys62; S-S1 in Fig. 2b) and C-terminus (Cys395 and Cys405; S-S2 in Fig. 2b)^10^.

**Figure 2.**
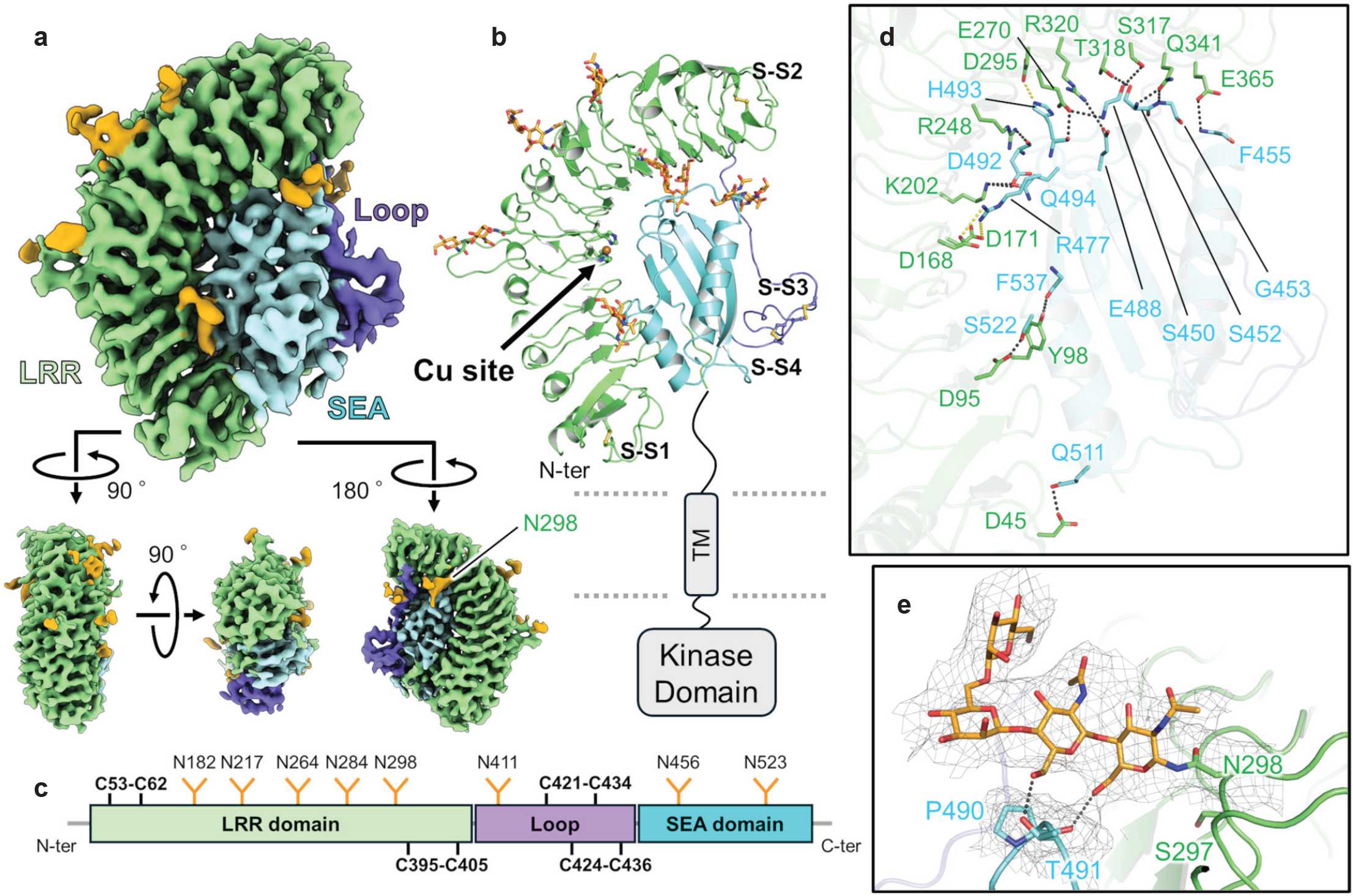
Cryo-EM structure of EctoCARD1. **a**, Overall structure of EctoCARD1 shown in four different orientations based on the cryo-EM map. **b**, Structural model of EctoCARD1, highlighting *N*-glycans, disulfide bonds, and the histidine residues involved in copper binding (shown as stick models). The copper atom represents a brown sphere. **c**, Schematic of EctoCARD1 domain architecture, with glycosylation sites, disulfide bond sites, and corresponding residue numbers indicated. **d**, Molecular interface between the LRR and SEA-like domains with possible hydrogen bonds and salt bridges depicted as black and yellow dotted lines, respectively. **e**, *N*-glycosylation at Asn298, showing possible hydrogen bonds between the sugar chain and the protein (black dashed lines). Cryo-EM maps are shown as gray mesh surfaces.

The role of glycosylation in LRR-RLKs has been well-documented, with notable examples including the tomato Cf-9 receptor^11^ and immune receptors in *Arabidopsis*^12^ and recently for MIK2 receptor^13,14^. Similarly, the cryo-EM map shows that at least eight asparagine residues in EctoCARD1 including Asn298 are modified by *N*-glycans. Notably, the *card1-7* allele, which carries a mutation at the neighboring Ser297, fails to exhibit the DMBQ-induced calcium elevation phenotype^4^. This impairment may be linked to disruptions in local glycosylation patterns on the CARD1 protein, which could compromise its function. Indeed, the carbohydrate chain to Asn298 interacts with the SEA-like domain (Fig. 2e), indicating that the glycosylation at this site plays an important role in retaining and stabilizing the EctoCARD1 structure.

### Cys421-Cys434 and Cys424-Cys436 pairs are not required for signaling

The reactivity of cysteine residues is influenced by two key factors: (i) the accessibility of the cysteine residue within the protein structure, and (ii) the local electrostatic environment surrounding the cysteine, including factors such as local pH. Higher pH (more alkaline), for instance, favors the thiolate anion form, thereby increasing susceptibility to oxidation^15–18^. Previous studies have suggested that specific cysteine residues play roles in sensing DMBQ (Cys395 and Cys405)^4^ and H_2_O_2_ (Cys421, Cys424, Cys434, and Cys436)^5^.

We thus investigated the physical properties of these cysteines within the EctoCARD1 structure. Our structural analysis reveals that Cys395 and Cys405 cysteines are located in the C-terminal end of the LRR solenoid structure, while Cys421, Cys424, Cys434, and Cys436 are in the loop region of EctoCARD1 protein (Fig. 3a). Importantly, these cysteines have formed disulfide bonds. The disulfide bond between Cys395 and Cys405 likely forms a C-terminal cysteine-pair capping domain – a feature commonly seen in plant LRRs. Cysteine-pair capping domains are often associated with protein stability and function (Fig. 3a)^10,19^. To examine the role of Cys395 and Cys405 in CARD1 signaling, we generated *card1* complemented lines with C395S/C405S substitutions (CARD1^C395S/C405S^). These mutants exhibited varying protein levels (Supplemental Fig. 4a). Similar to the single mutants of Cys395 and Cys405, CARD1^C395S/C405S^ mutant showed additional bands on protein blots, suggesting possible protein degradation^4^ (Supplemental Fig. 4a). Transformants with higher protein expression exhibited weak responses to both DMBQ and H_2_O_2_ compared to WT-AEQ (Fig. 3b-d), suggesting that Cys395 and Cys405 substitution may disrupt CARD1 structural integrity. This would in turn reduce the ability of CARD1^C395S/C405S^ to perceive quinones and H_2_O_2_, while increasing its susceptibility to endogenous proteases, leading to the accumulation of the truncated CARD1^C395S/C405S^.

**Figure 3.**
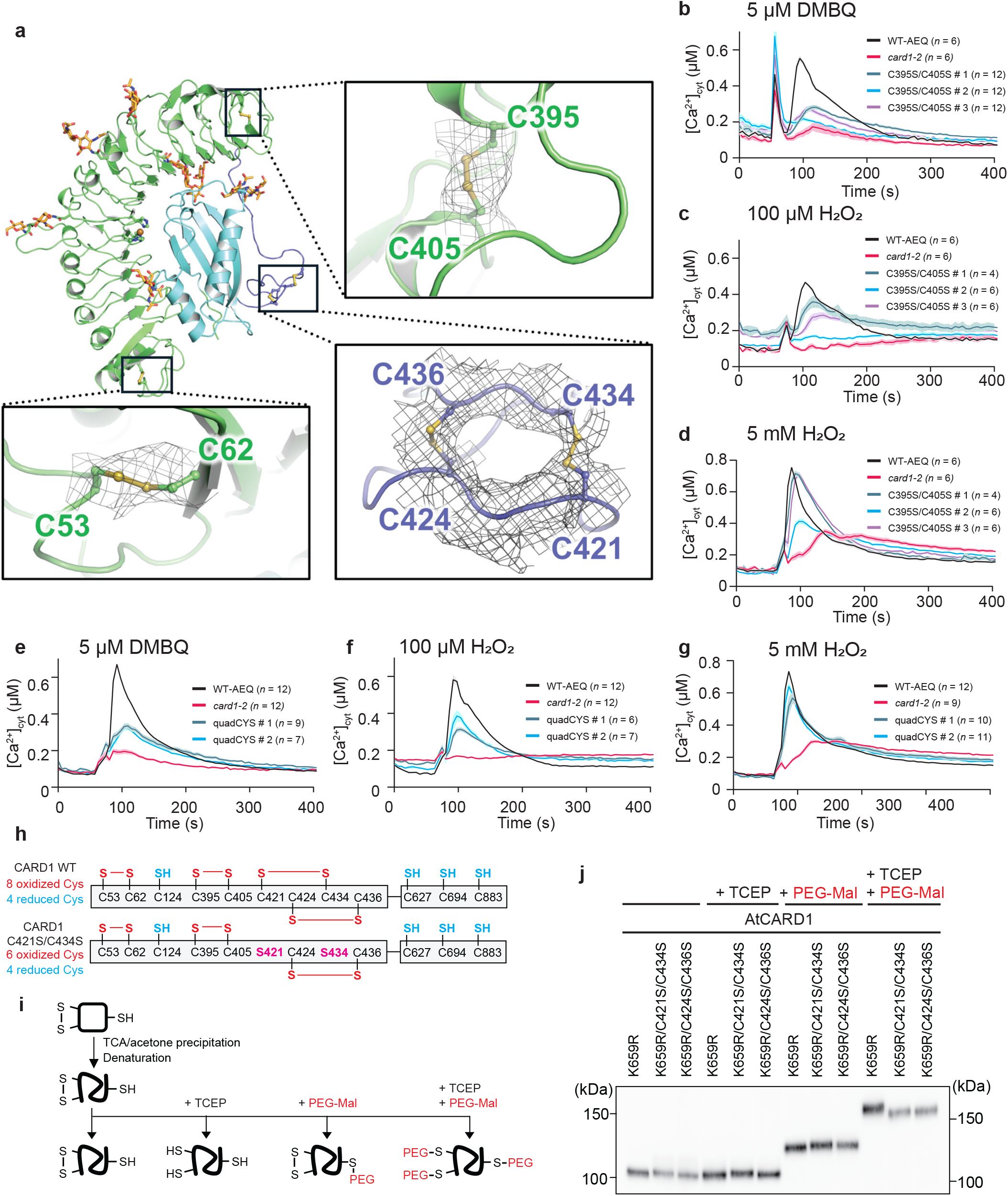
Disulfide bonding between Cys421-Cys434 and Cys424-Cys436 are dispensable for CARD1 H_2_O_2_ signaling. **a**, Structure model of EctoCARD1, highlighting disulfide bonds between cysteines pairs at the N- and C-terminal end of the LRR domain (green) and between cysteines in the loop region (blue). Cryo-EM maps surrounding these disulfide bonds are shown as gray mesh surfaces. **b-d**, [Ca^2+^]_cyt_ response of *Arabidopsis* seedlings WT-AEQ, *card1-2* and three independent transformants of *card1-2/CARD1*^*C39S5/C405S*^ in response to (**b**) DMBQ, (**c**) 100 µM H_2_O_2_ and (**d**) 5 mM H_2_O_2_. Stimuli were applied at 60 s. Data are mean ± SE with the number of seedlings indicated in each panel). **e-g**, [Ca^2+^]_cyt_ response of *Arabidopsis* seedlings WT-AEQ, *card1-2* and 2 independent transformants of *card1-2/CARD1*^*quadCYS*^ in response to (**e**) DMBQ, (**f**) 100 µM H_2_O_2_ and (**g**) 5 mM H_2_O_2_. Stimuli were applied at 60 s. Data are mean ± SE (number of seedlings are indicated in each panel). **h**, Possible redox state of cysteine residues in CARD1 and cysteine mutants, as predicted based on the determined EctoCARD1 structure and the CARD1 kinase domain structure predicted by Alphafold. **i**, Schematic of cysteine labeling workflow using PEG-Mal labeling. **j**, CARD1 and cysteine mutant migration patterns after PEG-Mal labeling. Protein extracts from *N. benthamiana* leaves expressing a kinase-dead AtCARD1 protein (CARD1^K659R^), or in combination with cysteine mutations (C421S/C434S and C424S/C436S). Protein blot was probed with anti-CARD1 antibody.

Next, we investigated the role of Cys421, Cys424, Cys434, and Cys436 in quinone and ROS perception. In the EctoCARD1 structure, all four cysteines form disulfide bonds (Cys421-Cys434 (S-S3) and Cys424-Cys436 (S-S4); Figs. 2b, 3a). This disulfide bonding pattern is intriguing as it was previously suggested that the formation of disulfide bonds between Cys421-Cys424, and Cys434-Cys436 is important for ROS sensing^5^. Our structural analysis revealed that the distance between Cys421 and Cys434 is more favorable for bond formation than between Cys421 and Cys424, although one cannot rule out that disulfide bonding may be exchanged between cysteines during ROS perception. To clarify the role of these cysteine residues, we initially checked their involvement in H_2_O_2_ signaling in the single cysteine mutants. However, no effect was observed on ROS perception at either low (100 µM) or high (5 mM) concentrations (Supplemental Fig. 4b, c). We then generated a quadruple cysteine mutant in *card1-2* background (*card1-2/pCARD1:CARD1*^*quadCYS*^) in which Cys421, Cys424, Cys434, and Cys436 were mutated to serine, and tested its ability to complement Ca^2+^ signaling phenotype in response to H_2_O_2_ or DMBQ. The CARD1^quadCYS^-expressing plants responded to quinone and low H_2_O_2_ (100 µM) stimuli, albeit at a slight reduction in Ca^2+^ bursts compared to WT-AEQ plants (Fig. 3e, f). Since the CARD1^quadCYS^ protein also displayed a faint additional band on the blot, it is likely that the protein stability is affected, although not as much as the CARD1^C395S/C405S^ protein (Supplemental Fig. 4a, d). At high H_2_O_2_ concentration (5 mM), Ca^2+^ signaling was similar between the transformants and WT-AEQ (Fig. 3g). Overall, genetic and structural data suggest that Cys421, Cys424, Cys434, and Cys436 were unlikely to play an important role in CARD1 signaling upon the perception of quinone and H_2_O_2_.

Our structural analysis reveals that EctoCARD1 contains one reduced cysteine (Cys124) and four disulfide bonds (Cys53-Cys62, Cys395-Cys405, Cys421-Cys434, and Cys424-Cys436) (Fig. 3a). The predicted structure of CARD1 kinase domain by AlphaFold contains 3 reduced cysteine residues (Cys627, Cys694, and Cys883) (Supplemental Fig. 5a). Thus, CARD1 is estimated to have 4 reduced and 8 oxidized cysteine residues (Fig. 3h). To validate disulfide pairing in EctoCARD1 *in vivo*, we used a thiol-specific labelling approach using a 2 kDa polyethylene glycol-conjugated maleimide (PEG-Mal), which selectively reacts with free thiol groups and add a PEG moiety per modified cysteine (Fig. 3i). PEGylation induces an SDS-PAGE mobility shift proportional to the number of PEG molecules added^20^. Unlabeled native CARD1 from untreated *Arabidopsis* showed a band at estimated molecular weight (102 kDa) while PEGylated CARD1 shifted to higher molecular weight (120 kDa) (Supplemental Fig. 5b). In *N. benthamiana*, we used the kinase-dead mutant of *Arabidopssis* CARD1 (CARD1^K659R^) that does not induce cell death phenotype associated with CARD1 overaccumulation (Supplemental Fig. 5c) and the protein migrated similarly to native CARD1 from *Arabidopsis* (Supplemental Fig. 5b). Protein bands from CARD1^K659R^ with substitutions of Cys421 and Cys434 to Ser (CARD1^K659R/C421S/C434S^) or Cys424 and Cys436 to Ser (CARD1^K659R/C424S/C436S^) showed comparable migration to that of CARD1^K659R^ in unlabeled samples (Fig. 3j). However, in PEG-Mal labeling samples after TCEP reduction (+ TCEP + PEG-Mal), both PEGylated CARD1^K659R/C421S/C434S^ and CARD1^K659R/C424S/C436S^ migrate further than PEGylated CARD1^K659R^, confirming that PEG-Mal labeling assay provides sufficient resolution to distinguish one disulfide bond formation (or reduction) in CARD1 (Fig. 3j). Importantly, in samples without TCEP reduction prior to PEG-Mal labeling (-TCEP + PEG-Mal), both PEGylated CARD1^K659R/C421S/C434S^ and CARD1^K659R/C424S/C436S^ migrate at a similar distance to CARD1^K659R^, implying that neither C421S/C434S nor C424S/C436S mutations influence the number of reduced cysteine residues in EctoCARD1 (Fig. 3j). Overall, these findings collectively support the idea that Cys421-Cys434 and Cys424-Cys436 pairs form disulfide bonds *in vivo*.

### Copper ion coordination by the histidine triad is essential for CARD1 signaling

Given that cysteines do not appear to play an essential role in CARD1 signaling, we carefully search for other structural features in EctoCARD1 that may explain how CARD1 can perceive ROS or quinones. Further inspection revealed a connected cryo-EM map between His199 and His222, indicating the presence of a metal ion (Fig. 4a). This possible metal-binding site, composed of histidine residues His197, His199, and His222 (forming a His-triad), appears to adopt a slightly distorted T-shaped coordination geometry (Fig. 4a). T-shaped metal sites involving histidine residues are commonly found in copper-containing proteins such as lytic polysaccharide monooxygenase (LPMO)^21^, copper amine oxidase^22^, and peptidylglycine α-hydroxylating monooxygenase ^23^, which utilize dioxygen (O_2_) or H_2_O_2_ to perform enzymatic reactions.

**Figure 4.**
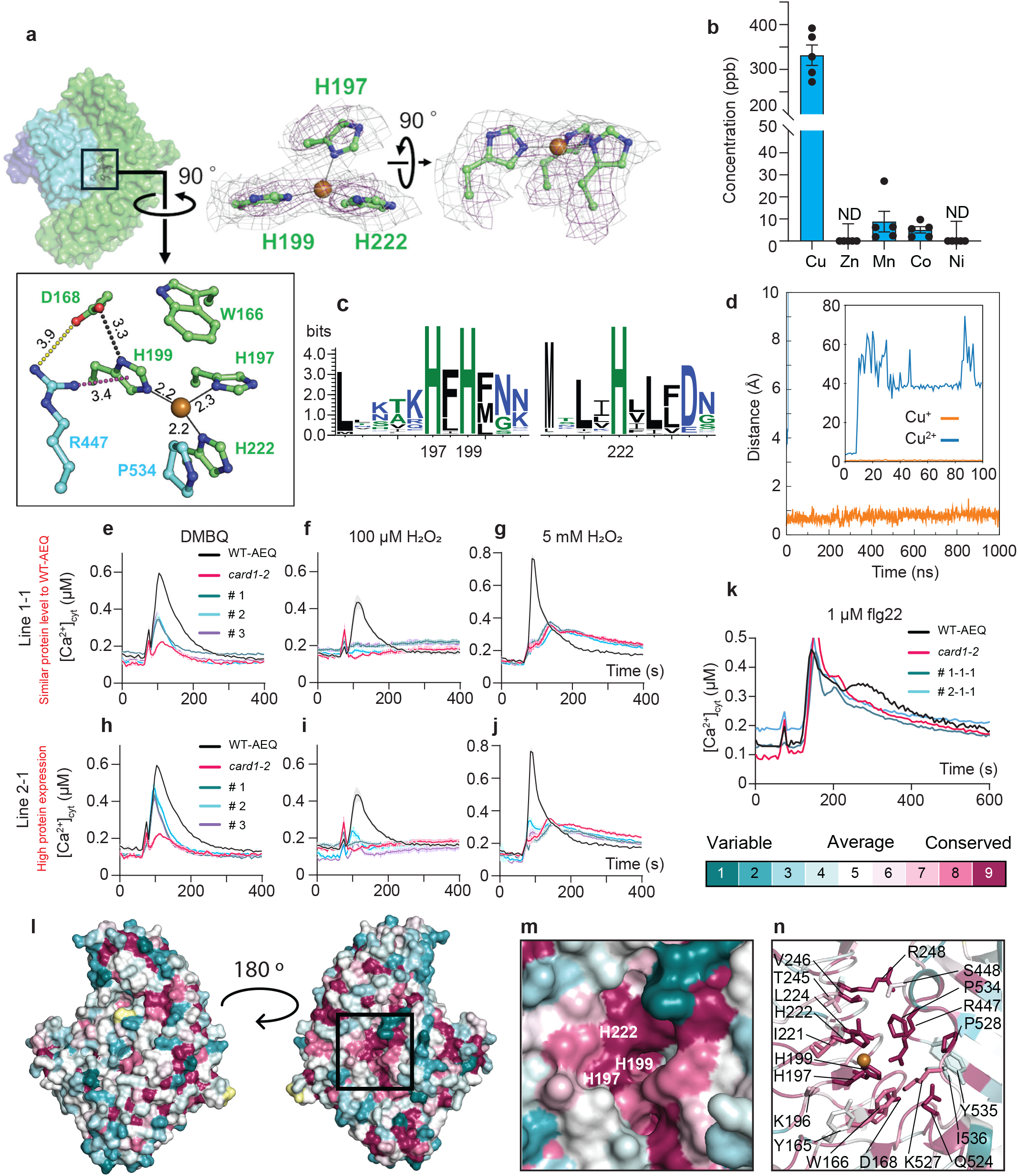
The His-triad in CARD1 protein is essential for proper quinone and ROS perception. **a**, Structure model of EctoCARD1 showing the copper (Cu)-binding site. Cryo-EM maps are shown at high and low thresholds (purple and gray meshes, respectively). The Cu-binding site is shown in two orientations. The inset provides a close-up view of the Cu-binding site, with copper depicted as a brown sphere. **b**, Quantification of metal elements bound to CARD1 by ICP-MS. **c**, Sequence logos generated with WebLogo server, showing the conservation of histidine residues in His-triad of land plants CARD1/CADLs. **d**, Time evolution of the distance between the copper ion (either Cu^+^ or Cu^2+^) and the geometric center of the His-triad (His197, His199, and His222) in the CARD1 during MD simulations. The Cu^2+^ ion rapidly dissociates from the His-triad shortly after the simulation begins. In contrast, the Cu^+^ ion remains stably coordinated to the triad throughout the 1-μs simulation, maintaining an average distance of 0.76 Å with a standard deviation of 0.17 Å. **e-g**, [Ca^2+^]_cyt_ response of *Arabidopsis* seedlings WT-AEQ, *card1-2*, and T3 lines (line 1-1-1 to line 1-1-3) of *card1-2/CARD1*^*HtriA*^ in response to (**d**) DMBQ, (**e**) 100 µM H_2_O_2_ and (**f**) 5 mM H_2_O_2_. Stimuli were applied at 60 s. For DMBQ treatment, data are mean ± SE (*n* = 6 for WT and *card1-2, n* = 12 for transformant lines. For H_2_O_2_ treatments, *n* = 6 for all genotypes. **h-j**, [Ca^2+^]_cyt_ response of *Arabidopsis* seedlings WT-AEQ, *card1-2*, and T3 lines (line 2-1-1 to line 2-1-3) of *card1-2/CARD1*^*HtriA*^ in response to (**g**) DMBQ, (**h**) 100 µM H_2_O_2_, and (**i**) 5 mM H_2_O_2_. Stimuli were applied at 60 s. For DMBQ treatment, data are mean ± SE (*n* = 6 for WT and *card1-2, n* = 12 for transformant lines. For H_2_O_2_ treatments, *n* = 6 for all genotypes. **k**, [Ca^2+^]_cyt_ response of *Arabidopsis* seedlings WT-AEQ, *card1-2*, and T3 lines of *card1-2/CARD1*^*HtriA*^ in response to 1 µM flg22. Data are mean ± SE (*n* = 6) **l**, Sequence conservation of CARD1 across angiosperm. Conservation scores of 19 CARD1 from angiosperms are calculated by the CONSURF program and mapped onto the EctoCARD1 structure. **m**, Surface view of the His-triad region (highlighted box in (**l**)). Histidine residues forming the His-triad are indicated. **n**, Ribbon diagram of the His-triad region. The side chains around the His-triad are shown in stick representation.

The metal ion is positioned on the plane formed by three ligand nitrogen atoms from the His-triad (Fig. 4a). The cryo-EM map around the His-triad indicates that His197 is more flexible than the other two His ligands, potentially flipping out and altering the coordination geometry of the metal site to a linear configuration. Additionally, His199 forms a hydrogen bond with nearby Asp168, which electrostatically interacts with Arg447 in the SEA-like domain. Arg447 further engaged in a cation-π interaction with His199’s imidazole ring. Further, His122 and Pro534 in the SEA-like domain show a van der Waals contact as is Trp166 with His197 and His199 (Fig. 4a). The surface of EctoCARD1 forms an exposed pocket that could accommodate small redox-active molecules, such as quinones or H_2_O_2_ (Supplemental Fig. 6a). To confirm that the copper ions are coordinated by the His-triad, we performed inductively coupled plasma mass spectrometry on purified EctoCARD1 protein. The results consistently showed copper as the most abundant metal in the sample (Fig. 4b, Supplemental Fig. 6b, c). We also confirmed that histidine residues within the triad are highly conserved in LRR-RLK Group VIII-1 (Fig. 4c) and it is absent in other LRR-RLK groups (Supplemental Fig. 6d). Altogether, these data indicate that copper is the primary metal which is coordinated by surface exposed His-triad that is highly conserved in CARD1.

The T-shaped geometry and the copper position on the plane formed by ligand N atoms imply that the Cu ion is likely in the reduced (Cu^+^) state, which may be oxidized to Cu^2+^ by H_2_O_2_. To characterize the binding affinity of a Cu ion to the CARD1, we performed molecular dynamics (MD) simulations using the cryo-EM structure of EctoCARD1. In these simulations, a Cu ion was manually placed in the His-triad, and the MD simulations were conducted for both Cu^+^ and Cu^2+^ binding to EctoCARD1. The results revealed that the Cu^+^ ion remained stably bound to the His-triad throughout the extended 1 μs simulation period, whereas the Cu^2+^ ion rapidly dissociated from the binding site upon simulation initiation (Fig. 4d, Supplementary Movie 1, 2). Time for Cu^+^ and Cu^2+^ localization within the His-triad was quantified and Cu^+^ significantly stayed within the His-triad in a longer arbitrary timeframe (Fig. 4d). These findings indicate that Cu^+^ binds stably to CARD1, whereas Cu^2+^ binding is unstable. The MD simulations strongly support the role of the His-triad as the Cu^+^ binding site, consistent with experimental observations and the coordination geometry reported in other copper-binding proteins^24,25^.

To validate the functional importance of the His-triad and Cu^+^ in CARD1, copper-free CARD1 mutant was generated by substituting histidine residues in His-triad to alanine residues (H197A/H199A/H222A; HtriA). We first experimentally determined the structure of EctoCARD1^HtriA^ by cryo-EM (PDB 9VBV) at a resolution of 3.61 Å (Supplemental Fig. 7, Supplementary Table 1). Superimposing the structures of EctoCARD1 and EctoCARD1^HtriA^ revealed no significant structural differences between the two proteins (Supplemental Fig. 8), suggesting that HtriA substitution did not affect the overall structure. Notably, a clear absence of density at the Cu-binding site in the cryo-EM map of EctoCARD1^HtriA^, compared to the EctoCARD1^WT^, confirms that the mutant protein is devoid of copper (Supplemental Fig. 8b, c). We then checked whether CARD1^HtriA^ could complement the stimuli-induced calcium elevation phenotypes in *card1-2* plants. The His-triad mutant (*card1-2/pCARD1:CARD1*^*HtriA*^) exhibited a severely impaired Ca^2+^ peak response to DMBQ in all 5 independent lines, irrespective of protein levels (Supplemental Fig. 9a), suggesting that the His-triad mutation significantly impairs DMBQ signaling (Fig. 4e, h, Supplemental Fig. fig. 9b). Importantly, the His-triad mutants were not able to response to low (100 µM) or high (5 mM) H_2_O_2_ as they phenocopied *card1-2* in their response to H_2_O_2_, even in transformants with high protein expression (Fig. 4f, g, i, j, Supplemental Fig. 9c, d). Additionally, the mutants also phenocopied *card1-2* in response to flg22 (Fig. 4k, Supplemental Fig. 9e). Overall, these data collectively imply that the His-triad, and thus Cu^+^ coordination, is crucial for signaling in CARD1, particularly for H_2_O_2_ perception.

To identify evolutionary conserved regions of CARD1 across angiosperms, we mapped sequence conservation onto the EctoCARD1 structure (Fig. 4l). This analysis revealed that not only the His-triad, but also surrounding residues, are highly conserved (Fig. 4m, n), implying that this region is crucial for CARD1 function and has been slowly evolving in CARD1 homologs. Altogether, these data further support the importance of Cu^+^ ion coordination by the His-triad in CARD1 for H_2_O_2_ perception.

## Discussion

CARD1 perceives quinone and H_2_O_2_ in plants and its homologs are widespread across the land plant lineage. Notably, multiple CARD1 homologs were identified in early-diverging land plants, such as the liverwort *Marchantia polymorpha* and the moss *Physcomitrium patens*. While gene family reduction is common in early land plants, the number of *CARD1/CADLs* homologs remains relatively unchanged, with eight in *Arabidopsis* and six in *M. polymorpha*. The functional significance of this family may depend on the expression patterns of individual homologs. For instance, *CADL3* – capable of perceiving both ROS and quinone (Fig. 1e, f) – is primarily expressed in the vasculature^26^. This suggests that certain CARD1 homologs may function in perceiving quinones or ROS produced in the vasculatures, potentially playing a role in vascular redox biology. Importantly, here we showed that apoplastic ROS generated by RBOHD are perceived by CARD1 during immune signaling (Fig. 1c, d). Consistent to this finding, both *card1* and *rbohd* mutants exhibit reduction in stomatal immunity^27^. Interestingly, both CARD1 and NADPH oxidases appear to have emerged concurrently in early land plants^28^. This co-evolution suggests that apoplastic ROS production and perception evolved together – a necessary adaptation as plants transitioned to land where ROS can no longer diffuses in the aqueous environment and required specific perception mechanisms.

The cryo-EM structure of plant-expressed EctoCARD1 revealed several unique features. First, *N*-glycosylation at Asn298 stabilizes intramolecular interaction between LRR domain and SEA domain-like fold (Fig. 2e). Second, the ectodomain of CARD1 features a cysteine-pair capping domain at both the N- and C-termini of the LRR domain, with an overall horseshoe shape typical of LRR-RLKs. However, it lacks an island domain, similar to other LRR-RLKs such as FLS2, PEPR and HAESA^3^. Instead, structural homology modelling using the DALI and Foldseek servers^29,30^ revealed that the C-terminal end of EctoCARD1 shares significant similarity with a SEA domain-containing proteins (Supplementary Tables 2, 3). In animals, SEA domains are commonly found in various cell surface receptors and often exhibit autoproteolytic activity^9,31^. The bacterial SEA-like domain is also involved in regulating intramembrane proteolysis *via* its autoproteolytic activity^32^. Notably, SEA domains are typically located near the C-terminal end of the ectodomain, just before the transmembrane domain^31^, which is also the case for CARD1 SEA domain-like fold. It is tempting to speculate that the SEA domain-like fold of CARD1 may undergo autoproteolysis, similar to other known mammalian SEA-domain containing receptors, thus modulating protein function *in vivo*.

Our analyses unequivocally demonstrate that Cys421, Cys424, Cys434, and Cys436 in CARD1 are not essential for signaling function. Further, these cysteine residues form disulfide bonds between Cys421-Cys434 and Cys424-Cys436, rather than between Cys421-Cys424 and Cys434-Cys436 as previously proposed^5^. In addition, PEG-Mal assay revealed disulfide bonding between Cys421-Cys434 and Cys424-Cys436 in unstimulated plants, suggesting that these disulfide bonding are redox-insensitive (Fig. 3j, Supplemental Fig. 5b). These disulfide bonds in the exposed loop region (Figs. 2a, 3a) may play a role in protecting CARD1 from proteolytic cleavage by endogenous proteases.

We identified a novel His-triad in the CARD1 ectodomain, marking the first instance in plant LRR-RLKs shown to coordinate metal ions (Cu^+^), that is crucial for H_2_O_2_ perception as the loss of Cu^+^ in CARD1^HtriA^ showed no H_2_O_2_-induced Ca^2+^ signaling (Fig. 4e-k). The conservation of histidine residues forming Cu^+^ binding site across angiosperms supports the importance of Cu^+^ for CARD1 function (Fig. 4c, m). While copper is also involved in ethylene receptor function^33^, the role of copper in CARD1 is distinct. Many plant redox regulation systems or sensors rely on cysteine-based redox couples or methionine-based oxidative post-translational modifications^15–18^. The apoplastic environment, that is unique to plant cells and is more oxidized than the cytosol, may require distinct mechanisms for sensing redox-active molecules^17^. The Cu^+^-bound EctoCARD1 structure determined in this study likely represents an unstimulated state, as neither H_2_O_2_ nor quinones were added during sample preparation prior to structural determination. Under aerobic conditions in the apoplast, Cu^+^ is expected to be spontaneously oxidized to Cu^2+^, suggesting that a mechanism must exist to maintain extracellular Cu ions in the reduced Cu^+^ state. A potential source of reducing power for this is ascorbic acid which accumulates in the apoplast at concentrations on the order of 0.1 mM^34^ and is capable of reducing Cu^2+^ to Cu^+^. Alternatively, plants may employ the ferric reductase oxidases FRO4 or FRO5 to mediate the reduction of Cu^2+^ to Cu^+^ in the apoplast^35^.

We have successfully determined how CARD1 uses Cu^+^ in CARD1 to perceive ROS and therefore demonstrate that CARD1 can discriminate between quinone and H_2_O_2_. However, the exact mechanism by which CARD1 distinctively perceives between quinone from H_2_O_2_ signals remains to be investigated. We cannot exclude the possibility that oxidation of Cu^+^ by H_2_O_2_ results in the release of Cu ion from the His-triad. However, it is unlikely to be sufficient as a trigger of CARD1 activation since the elimination of the Cu ion did not significantly affect the EctoCARD1 structure (Supplemental Fig. 8). Based on this data, we propose that CARD1 may act as an H_2_O_2_ signal modifier *via* its catalytic Cu^+^ and suggest two possible models by which Cu^+^ in CARD1 could mediate H_2_O_2_ signaling (Supplemental Fig. 10). In one model, catalytic Cu^+^ facilitates the generation of secondary signaling molecules, such as quinones, through the generation of Fenton reaction-derived hydroxyl radicals (Supplemental Fig. 10; off-Cu model). In an alternative model, signal generation may proceed *via* reactive copper-oxyl (^•^O-Cu^2+^) intermediates (Supplemental Fig. 10; on-Cu model). Indeed, Cu ions are well known to function as cofactors for various metalloenzymes^36^. For example, LPMOs are copper-dependent enzymes that catalyze the oxidative cleavage of polysaccharide. Notably, LPMO utilizes H_2_O_2_ as a co-substrate, with Cu^+^-mediated homolytic cleavage of H_2_O_2_ being a key step in this enzymatic reaction^37^. Similarly, CARD1 may also use H_2_O_2_ as a co-substrate to generate quinones in the presence of a specific substrate, thereby activating CARD1 signaling pathway. Overall, our work provides new insights into how plasma membrane receptors can integrate metal ions for signal perception and lays the groundwork for further research into ROS sensing mechanisms in plants.

## Acknowledgement

We thank Mikiko Fukatsu, Yoko Nagai, and the ITbM office for supports; Shirasu groups and the ITbM members for various discussions; Mr. Hiroki Tanino (Graduate School of Pharmaceutical Sciences, the University of Osaka) for support in cryo-EM data collection.

## Funding

Japan Society for the Promotion of Science KAKENHI grant JP22H00364 (to K.S)

Japan Society for the Promotion of Science KAKENHI grant JP24K01718 (to N.I, Y.F, A.L)

Japan Science and Technology Agency (JST) GteX program grant JPMJGX23B2 (to K.S)

Japan Science and Technology Agency (JST) FOREST program grant JPMJFR220G (to A.L)

MEXT Promotion of Development of a Joint Usage/Research System Project: Coalition of Universities for Research Excellence Program (CURE) grant JPMXP1323015482 (to K.J.F, Y.T)

MEXT Project for promoting public utilization of advanced research infrastructure (Program for supporting construction of core facilities) Grant Number JPMXS04411024 (to H.I, M.N)

RIKEN TRIP initiative Field Omics (to K.S)

Mitsubishi Foundation (to A.L)

## Contributions

Conceptualization, NI, YF, KS, AL; Methodology, NI, YF, KJF, HI, MN, TY, TI, KS, AL; Investigation, validation and analysis, NI, YF, YS, KT, RH, KJF, HI, MN, AL; Visualization, NI, YF, KJF, HI, MN, KS, AL; Resources, NI, YF, KJF, TY, TI, KS, AL; Supervision, NI, YT, TI, KS, AL; Writing – original draft, NI, YF, KJF, AL; Writing – review and editing, NI, YF, KJF, MN, TY, TI, KS, AL with all authors’ inputs.

