## Supplemental information for "A copper-dependent, redox-based hydrogen peroxide perception in plants"

**Supplemental information for Ishihama et al.,**

#### Methods

**Chemical and Reagents.** Chemicals were purchased from Fujifilm Wako Pure Chemical (Fujifilm Wako Pure Chemical Cooperation) unless otherwise mentioned. The flg22 peptide (QRLSTGSRINSAKDDAAGLQIA, 95.3% purity) was custom-synthesized by Medical & Biological Laboratory, and dissolved in endotoxin-free water.

**Plant materials.** All *A. thaliana* lines are Col-0 background. Transgenic *Arabidopsis* lines expressing (apo)aequorin (denote as WT-AEQ), *card1-2*, complemented *CARD1* and *CARD1* with single cysteine mutations were previously described[4]. Seeds of *rbohd* and *rbohf* were previous published[38]. Seeds were surface-sterilized and stratified for 2 d before growing in liquid medium (1/2 MS, 0.25% sucrose, pH 5.7) at 20-22 ºC, 16-h photoperiod, 60-80 µE. Soil grown *Arabidopsis* plants were grown individually in a 6 cm diameter pot (1:1 mixture of soil and vermiculite) under the same condition. For $Ca^{2+}$ measurement experiments, liquid grown *Arabidopsis* seedlings were grown for 1week post-germination. For experiments with soil-grown *Nicotiana benthamiana*, plants were grown for 4-5 weeks post-germination in an individual pot (10 cm diameter with 1:1 soil and vermiculite) under control conditions (25-26 ºC, 16-h photoperiod, 100-150 µE). Where possible, plants were randomly selected from the larger pools which were grown in identical conditions. Different genotypes were randomly positioned in the growth chambers/shelves.

**Cloning and transformation.** Primers used in this study are provided in Supplementary Table 4. All sequences were verified by Sanger sequencing. To generate pHREAC-EctoCARD1-MBP and pHREAC-EctoCARD1$^{HtriA}$-MBP for cryo-EM study, the CARD1 ectodomain (residues 1–546), containing a GSA linker and a codon-optimized maltose-binding protein (MBP) at its C-terminus (EctoCARD1-MBP), was cloned into the *Bsa*I site of pHREAC vector[39] using In-Fusion HD cloning kit. EctoCARD1 part was cloned from genomic DNA whilst codon-optimized MBP was synthesized (Eurofin). The plasmid was electroporated into *A. tumefaciens* C58C1 strains. See Supplmentary Data 1 for nucleotide sequence of EctoCARD1-MBP.

To generate pHREAC-AtCARD1$^{K659R}$, pHREAC-AtCARD1$^{K659R/C421S/C434S}$ or pHREAC-AtCARD1$^{K659R/C424S/C436S}$ for PEG-Mal labeling experiments in *N. benthamiana*, PCR primers were designed with 15 bp overlapping sequence at their 5' end. Cysteine-coding codons and lysine-coding codon were replaced with serine- and arginine-coding codons, respectively. PCR was performed with full-length *CARD1* cDNA sequence to yield mutagenized *CARD1* cDNA fragments using KOD one (TOYOBO). These fragments were combined with linearized pHREAC vector using In-Fusion HD cloning kit. The plasmid was electroporated into *A. tumefaciens* C58C1 strains

For complementation experiments in *Arabidopsis* with mutagenized *CARD1*, PCR primers were designed such that codons of targeted amino acid were replaced with codons of desirable amino acid changes. Further, primers were also designed with 15 bp overlapping sequence at their 5' end. PCR was performed with full-length *CARD1* gDNA sequence to yield mutagenized *CARD1* gDNA fragments using KOD one. Native *CARD1* promoter (2.7 kb 5′ regulatory sequences of *CARD1*; *pCARD1*) fragment was amplified from Col-0 gDNA.  Fragments were combined with either *Bsp*1407I-inearized pGWB4 binary vector[40] , or with *Bam*HI/*Xho*I-linearized pKI1.1R binary vector[41]. Since *Bam*HI/*Xho*I-linearized pKI1.1R binary vector eliminates Cas9 and guide RNA scaffold coding regions, the vector effectively allows for red fluorescent seed selections in plants. The *AtCADL1*, *AtCADL3*, *CARD1*$^{C39S/C40S5}$ and *CARD*$^{quadCYS}$ were cloned into *Bsp*1407I-inearized pGWB4 binary vector with 2 fragments In-Fusion Snap assembly cloning, while the *CARD1*$^{HtriA}$ was cloned into *Bam*HI/*Xho*I-linearized pKI1.1R vector with 4 fragments In-Fusion Snap assembly cloning.

**Protein expression and purification.** The expression of CARD1 ectodomain was performed according to the previously described method[8] with some modifications. *Agrobacterium* harboring pHREAC-EctoCARD1-MBP was grown overnight and suspended in 10 mM MES-NaOH (pH 5.6) and 10 mM $MgCl_2$ to $OD_{600}$ = 0.2. The suspension was infiltrated with a needleless syringe into leaves of soil-grown *N. benthamiana*. From three to five days-post-infiltration, infected leaves were excised and immersed in collection buffer (20 mM Tris-HCl (pH7.5), 0.01% (v/v) Tween 20, and 1 × cOmplete EDTA-free Protease Inhibitor Cocktail (Roche)). The collection buffer was infiltrated into the leaf tissue by reducing the pressure to 60 hPa for 10 min and then slowly releasing the pressure for 10 min. After removing buffer residues, the apoplastic fluid was collected by centrifugation twice (1,500 *g*, 10 min, 4 °C) and subjected to anion exchange chromatography using a HiTrap Q HP column (Cytiva). Proteins were eluted by a linear NaCl gradient (from 50 mM to 300 mM in 20 mM Tris-HCl (pH7.5) and 0.1% (v/v) CHAPS over 15 column volume). Fractions containing EctoCARD1-MBP were verified by immunoblot using anti-MBP antibody (M091-3; MBL). Peak fractions of EctoCARD1-MBP were pooled and concentrated using a Vivaspin 500 (10kDa MWCO; Sartorius), and further purified by size exclusion chromatography on a Superdex200 Increase 10/300 GL column (Cytiva) in a buffer containing 24.5 mM Tris-HCl (pH7.5) and 150 mM NaCl. Fractions containing EctoCARD1-MBP were verified by SDS-PAGE followed by Coomassie staining (InstantBlue; abcam), pooled, and concentrated further using Vivaspin 500 (10 kDa MWCO). Protein samples were flash-frozen in liquid nitrogen and stored at −80 °C, in aliquots, until required.

**Cryo-EM sample preparation.** The WT and mutant protein samples were diluted by the purification buffer to be ~ 0.2 mg/mL. A QUANTIFOIL Cu R1.2/1.3 200 mesh grid (Quantifoil Micro Tools GmbH) was glow discharged by Auto Fine Coater JEC-3000FC (JEOL) at 20 mA and 7 Pa for 20 sec. 3 μL of samples were applied on the grids. The grids were then blotted for 3 sec and rapidly cryocooled in liquid ethane by a Vitrobot Mark IV (Thermo Fischer Scientific) device. The sample chamber of Vitrobot was kept at 8 °C and 100% humidity.

**Cryo-EM data acquisition and processing.** All datasets were collected using CRYO ARM™ 200 (JEOL) equipped with a K3 direct electron detector (Gatan) controlled by SerialEM[42] at Graduate School of Pharmaceutical Sciences, Osaka University. All movies were collected at 60,000 magnification (pixel size: 0.83 Å) and the defocus range was from −0.7 to −2.2. Total electron exposure was 40 e−/Å2. Detailed conditions are shown in Supplementary Table 1.

Collected movies for the WT and the mutant samples were processed with cryoSPARC v3.2 and CryoSPARC v4.5.1, respectively[43,44]. Collected movies were imported and motion corrected. Contrast transfer function (CTF) was estimated by a patch CTF estimation job and particles were picked by using a Blob Picking job. Selected particles were extracted. 2D classification was performed to remove junk particles. Ab-initio map reconstruction was performed by particles selected by 2D classification. Several refinement jobs including CTF refinement and aberration correction[45] were performed to estimate accurate particle orientations and to obtain better cryo-EM maps. The detailed workflows are shown in Supplemental Figs. 3 and 7.

**Model building and validation.** An initial model of EctoCARD1 was predicted by ColabFold[46]. The model was manually fitted to the map on ChimeraX[47]. Fitted models were refined by Phenix Real-space Refinement[48]. Obtained models were further modified and carbohydrate chains were built manually in Coot[49]. Real-space refinement and manual model building were repeated. Final models were validated by MolProbity[50]. Models and cryo-EM maps were visualized by PyMOL (The PyMOL Molecular Graphics System, Version 2.4.0 Schrödinger, LLC.) and ChimeraX to draw figures.

**Whole-seedling aequorin luminometry.** The measurement of cytosolic free calcium ($[Ca^{2+}]_{cyt)}$) in response to stimuli was performed as previously described[4]. In brief, liquid-grown seedlings were individually placed in wells of white 96-well plates (Lumitrac flat bottom, Greiner Bio-One) with coelenterazine solution (Biosynth) overnight. The next day, luminescence values were recorded in each sample using either a $Tristar^2$ LB942 modular or a Tristar 3 multimode microplate readers (Berthold Technology). After baseline luminescence values were recorded for 60 s, 100 µl of stimuli was injected (2 times the final concentration). Luminescence values were recorded every 5 s for 500 s. Discharge solutions (10% (v/v) ethanol, 1 M $CaCl_2$) were injected and the luminescence was recorded for a further 60 s. Calibration to convert luminescence values to $[Ca^{2+}]_{cyt}$ was performed using the following previously published equation. Seedlings which were poorly grown and confirmed not to be genetically related to the genotype and/or possibly due to sterilisation method used, were omitted from study. Plants which did not expressed sufficient amount of aequorin, possibly due to silencing, were omitted from the study.

**CARD1 immunoblot.** To detect CARD1 protein in *Arabidopsis*, seedlings were harvested, snapped frozen in liquid $N_2$ and stored in –80 ºC. Samples were ground to a fine powder with zirconia beads (3-mm diameter) with either a FastPrep-24 or a FastPrep-5G sample shakers (MP Biomedicals). Proteins were extracted in extraction buffer (50 mM Tris-HCl (pH 7.5), 150 mM NaCl, 10% glycerol, 2 mM EDTA, 5 mM DTT, 1 × EDTA-free cOmplete protease inhibitor cocktail, 0.1% IGEPAL CA-630, 0.5 mM phenylmethane sulfonyl fluoride) by incubating the samples for 30 min at 4 ºC. Insoluble debris was pelleted by centrifugation (10 min, 4 ºC, 20,000 *g*). Equal amounts of protein were loaded on to an acrylamide gel and subjected to SDS-PAGE (Tris-Glycine-based). Proteins were then electroblotted onto a PVDF membrane using a trans-blot turbo transfer system (Bio-Rad) according to the manufacturer's instructions. Membranes were blocked overnight at 4 °C in TBS-T (20 mM Tris, 150 mM NaCl, 0.1% (v/v) Tween 20, pH 7.5) containing 5% (w/v) skimmed milk powder. To detect CARD1, we incubated the blot with an anti-CARD1 peptide antibody (1:5,000)[4] for 1 h at room temperature with shaking. Blots were then further incubated for 1 h with peroxidase-linked anti-rabbit IgG (1:10,000; NA934, GE Healthcare). Luminescence signal was detected using Clarity Max Western ECL substrate (Bio-rad) according to the manufacturer's instructions with either a LAS 4000 system or a LAS4010 system (GE Healthcare). Equal protein loading was verified by staining the detected membrane blot with Coomassie blue solution (0.1% (w/v) CBB R250 in 5:4:1 (v:v:v) mixture of $H_2O$, methanol, acetic acid).

**Protein expression in *Nicotiana benthamiana* and PEG-Mal labeling.** Selective labeling of the thiol groups was carried out as described[20] with minor modifications. Samples were obtained from either *Arabidopsis* seedlings or from *N. benthamiana* leaves transiently expressing AtCARD1 variants. Samples were ground into a fine powder with liquid nitrogen and then homogenized in 10% trichloroacetic acid in acetone. The homogenate was incubated at −20 °C for 1 h and centrifuged (20,000 *g* for 10 min at 4 °C). The supernatant was discarded, and the pellet was washed twice with ice-cold acetone. The pellet was resuspended in resuspension buffer (50 mM HEPES-NaOH (pH 7.0) and 2% SDS) containing either 10 mM 2k PEG-maleimide (2k PEG-Mal; SUNBRIGHT ME-020MA, NOF) or 40 mM tris (2-carboxyethyl) phosphine (TCEP; 203-20153, FUJIFILM Wako Pure Chemicals) for 90 and 30 min, respectively. After centrifugation (20,000 *g* for 10 min), the supernatant was collected. A 100 μL aliquot from the TCEP-treated sample was transferred to a new tube, and proteins in the sample were precipitated using chloroform/methanol[51]. The pellet was washed twice with ice-cold acetone and then resuspended with resuspension buffer containing 10 mM 2k PEG-Mal as described above. The samples were mixed with 2× SDS-PAGE sample buffer (without reducing agent) and subjected to immunoblot analysis.

**Inductively coupled plasma time-of-flight mass spectrometry (ICP-TOF-MS).** Determination of metal ions bound to EctoCARD1 was performed using the OptiMass 9600 (GBC Scientific Equipment Pty Ltd). The instrument was tuned using a 10 ng/mL multi-element standard solution (Agilent). The concentrations from 0.1 to 10 ng/mL of various standard metal solutions diluted in buffer solution (1 mM Tris-HCl, 10 mM NaCl, pH 7.0) were prepared separately by dilution of standard stock solutions (copper (cat. number 033-16201), zinc (cat. number 266-02341), manganese (cat. number 130-19461), cobalt (cat. number 035-26151) and nickel (cat. number 141-10151); all from FUJIFILM Wako Pure Chemical) with 2% $HNO_3$. The instrument was optimized and calibrated to measure the concentration of different metal ions individually. Optimized operating conditions are described in Supplementary Table 5. EctoCARD1 protein samples were acidified with 2% $HNO_3$ prior to analysis by ICP-TOF-MS. SEC buffer (24.5 mM Tris-HCl pH7.5, and 150 mM NaCl) were prepared in the same way as EctoCARD1 protein and analyzed by ICP-TOF-MS and used as control. Intensity of the control was subtracted from EctoCARD1 samples to obtain relative metal concentrations in EctoCARD1.

**Molecular Dynamic Simulations.** The stability of copper ion binding was assessed through molecular dynamics (MD) simulations. The cryo-EM-derived three-dimensional structure of EctoCARD1 was used as the target protein for the copper ion. Computational models for the MD simulations were created by manually positioning a copper ion ($Cu^+$ or $Cu^{2+}$) within the histidine triad composed of His197, His199, and His222 in the EctoCARD1 protein. These models were placed in a periodic boundary water box with dimensions of 65 × 97 × 112 Å[52]. The simulations ran for up to 1 μs with a 2 fs time step under constant pressure and temperature (NPT) conditions of 1 atm and 300 K, while maintaining a constant number of particles. Temperature was incrementally increased from 0 to 300 K and maintained at 300 K using a Langevin thermostat[52]. The particle mesh Ewald (PME) method[53] was used to take into account the non-bonding interactions, while the SHAKE method[54] enforced distance constraints on hydrogen-containing bonds. To maintain charge neutrality, nine and eight sodium ions were added in the $Cu^+$ and $Cu^{2+}$ systems, respectively. The periodic boundary box contained a total of 57,966 and 57,971 atoms for the $Cu^+$ and $Cu^{2+}$ systems, respectively. TIP3P[55] and ff14SB7[56] were used for water molecule and protein force fields, respectively. The Li/Merz monovalent ions force field[57] was applied to the $Cu^+$ ion, while the Li/Merz highly charged ions force field[58] was used for the $Cu^{2+}$ ion. All MD simulations were performed using the AMBER 2022 program package[59].

**CARD1 sequence mining and phylogenetic analysis and bioinformatics.** Sequences of CARD1 homologs from other plant species were obtained from Phytozome (database version 12.1) as previously described[4]. Briefly, full-length *Arabidopsis* CARD1 amino acid sequence (*At5g49760*) was used as a query for blastP against the database with the cut-off E-value of $1 \times 10^{-100}$ and hit sequences with $\geq$ 80% query coverage were selected. In addition to previously published *Phtheirospermum japonicum* and *Striga* CADL homologs[4], a total of 405 CADL sequences were obtained, spanning across 57 land plant species (Supplementary Data 2). Amino acid sequence alignment was performed on full-length sequences using MUSCLE (default parameters)[60] and the alignment were further processed using trimAl with 'Gappyout' automated trimming method[61]. A phylogenetic tree was inferred from a trimmed alignment with RAxML[62], using the PROTGAMMAAUTO model, and rapid bootstrapping with the autoMRE bootstrapping, using a random number seeds for a full maximum-likelihood tree search. The resulting estimated phylogeny was viewed as unrooted tree using iTOL[63]. His-triad conservation in land plant species were illustrated with Weblogo[64] For Consurf analysis, CARD1 homologs were identified from the NCBI non-redundant protein sequences database, using *Arabidopsis* CARD1 amino acid sequence, with a blastP search. Amino acid sequences of CARD1 homologs from 17 plant species (Supplementary Data 3) were aligned using Clustal Omega[65]. Conservation scores were calculated with ConSurf server[66] and mapped to the EctoCARD1 structure determined in this study.

**Data presentation, statistics and availability.** Data and graphs are presented and analyzed using GraphPad Prism 10 (Dotmatics) or Excel (Microsoft). All data in this study are available in the article or in supplementary information. Amino acid sequences used in this study are provided in Supplementary Data 2 and 3. The coordination files and cryo-EM maps have been deposited in the Protein Data Bank and the Electron Microscopy Data Bank (EMDB) under the accession codes PDB 9VBD/EMD-64920 (WT) and PDB 9VBV/EMD-64934 (HtriA). Other data are available from the corresponding author as request.

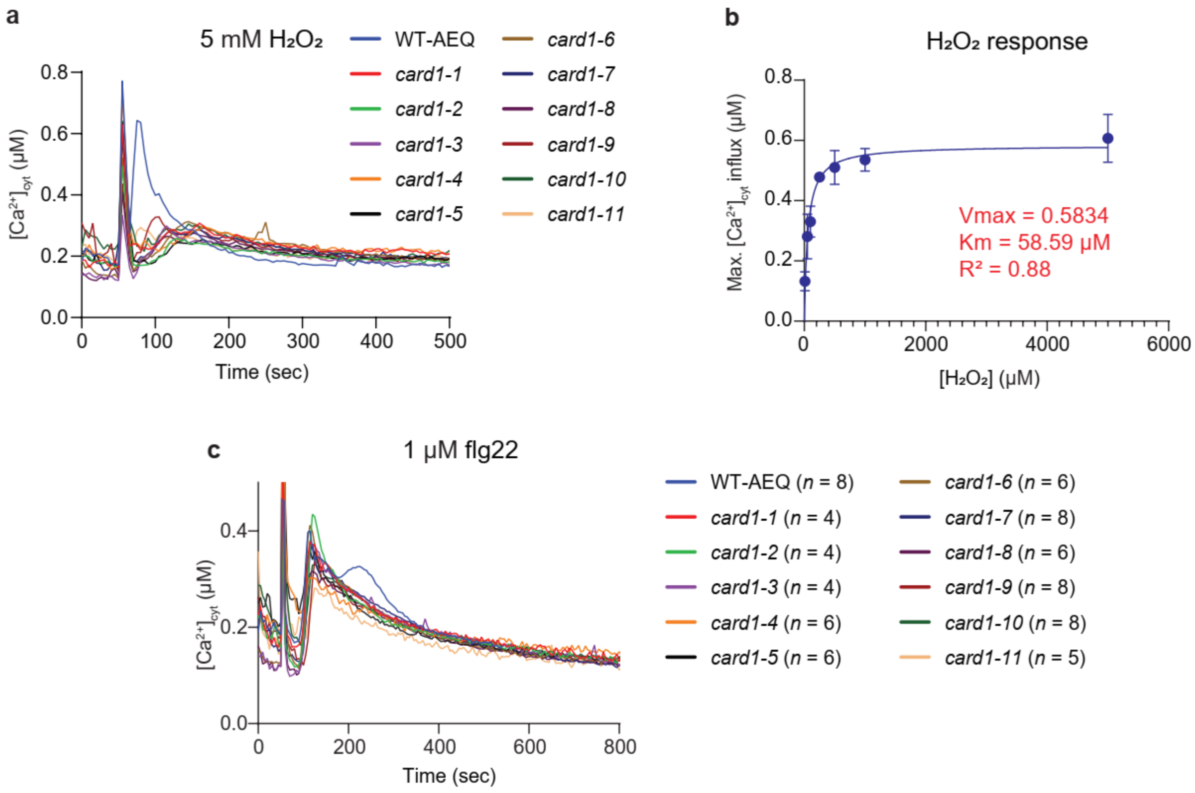

**Supplemental Figure 1. CARD1 is required for apoplastic ROS perception. a,** $H_2O_2$-induced $[Ca^{2+}]_{cyt}$ elevation is impaired in *card1* mutant alleles. Stimuli (5 mM $H_2O_2$) was applied at 60 s and data were measured every 5 seconds. Data are means, $n$ = 4 for all genotypes. Standard error (SE) was excluded for clarity. **b,** $H_2O_2$-induced $[Ca^{2+}]_{cyt}$ elevation follows standard Michaelis-Menten kinetics. Data are mean ± SE. $n$ = 6. **c,** flg22-induced $[Ca^{2+}]_{cyt}$ elevation showed two distinct peaks in wild-type (WT) response, but the second peak is absent in *card1* alleles. Stimuli (1 μM flg22) was applied at 60 s and data were measured every 5 seconds. Data are means; $n$ values are indicated for each genotype/alleles. SE was excluded for clarity.

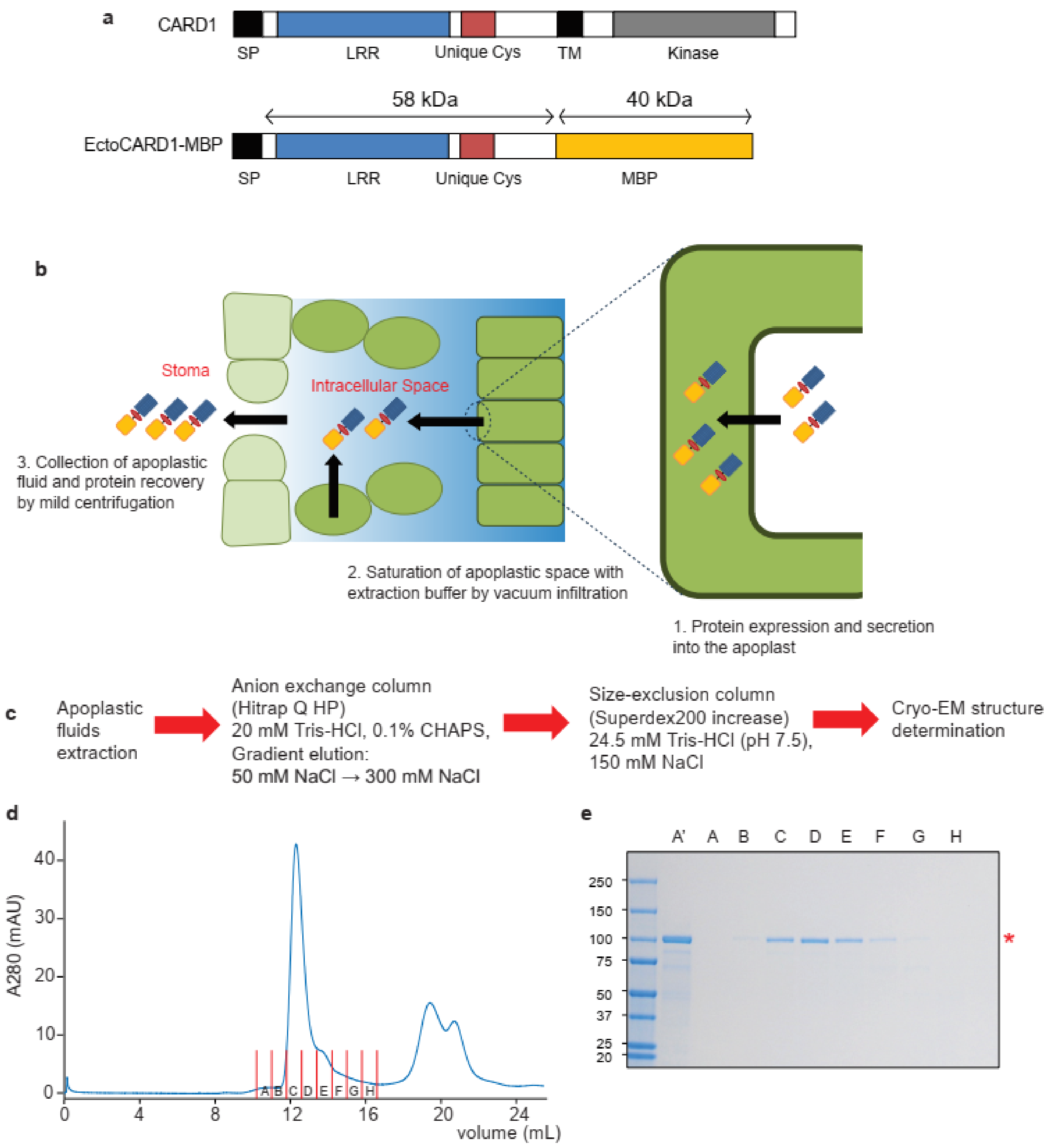

**Supplemental Figure 2. Expression and purification of EctoCARD1 from *Nicotiana benthamiana*. a,** Simplified diagram of the full-length CARD1 protein (top) and the EctoCARD1-MBP construct (bottom), highlighting different domains. SP, signal peptide; LRR, leucine-rich repeat; Unique Cys, a unique cluster of cysteine residues; TM, transmembrane; MBP, maltose-binding protein. **b,** Expression of EctoCARD1-MBP in *N. benthamiana* and its accumulation in apoplast. **c,** Strategic overview of the purification strategy for EctoCARD1-MBP prior to cryo-EM structure determination. **d,** Fast protein liquid chromatography trace of pooled fractions from anion exchange chromatography containing EctoCARD1, which were subsequently subject to size-exclusion chromatography (SEC). Fraction A to H were collected. **e,** Collected fractions were analyzed by SDS-PAGE. Fraction A' indicates the pooled sample from anion exchange chromatography. Fraction A to H correspond to SEC elution as shown in (**d**).

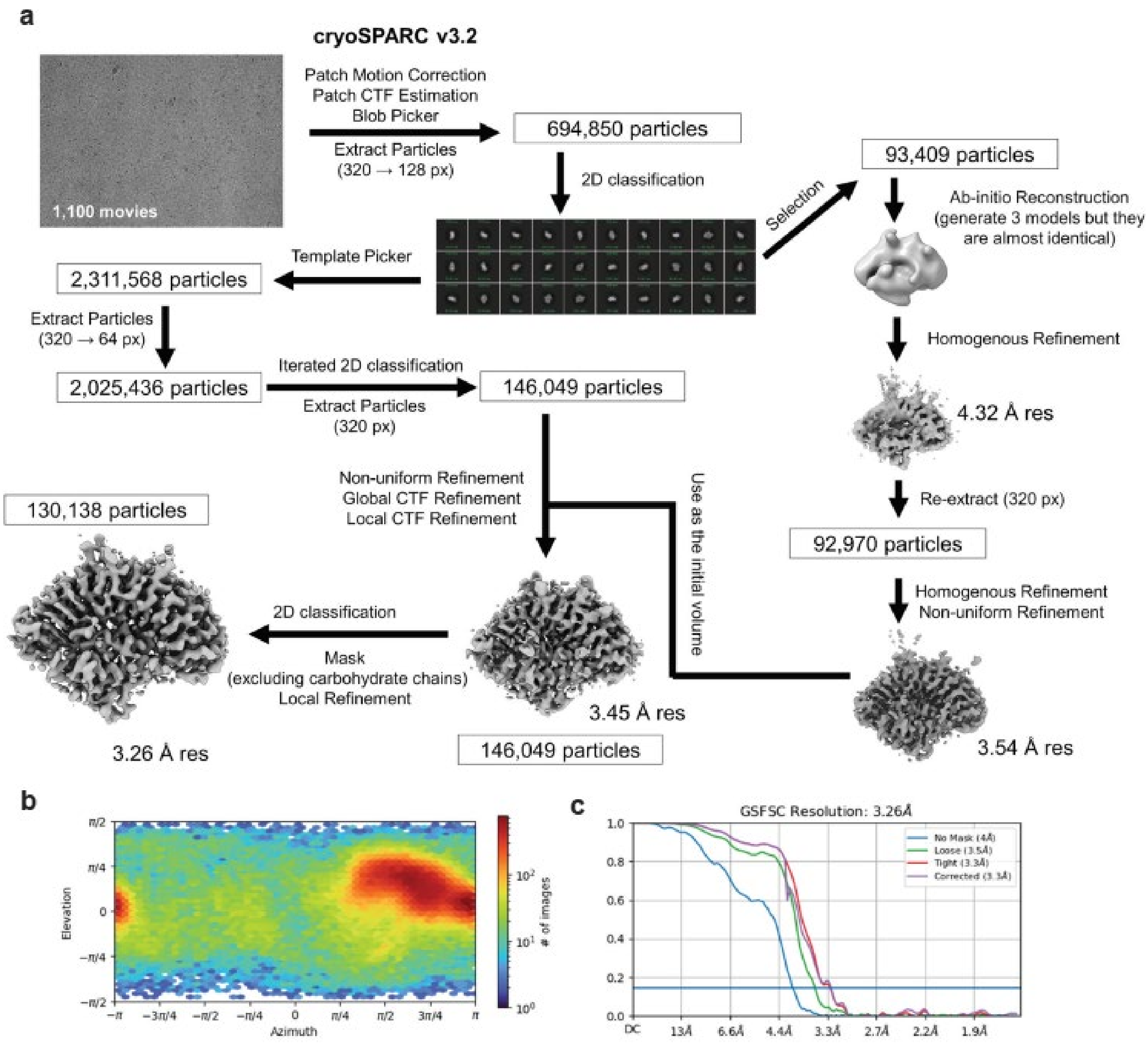

**Supplemental Figure 3. Cryo-EM data processing for WT EctoCARD1. a,** Workflow of data processing of WT EctoCARD1. **b,** Particle orientation distribution. **c,** Gold standard Fourier shell correlation (GSFSC) curves calculated by using different masks.

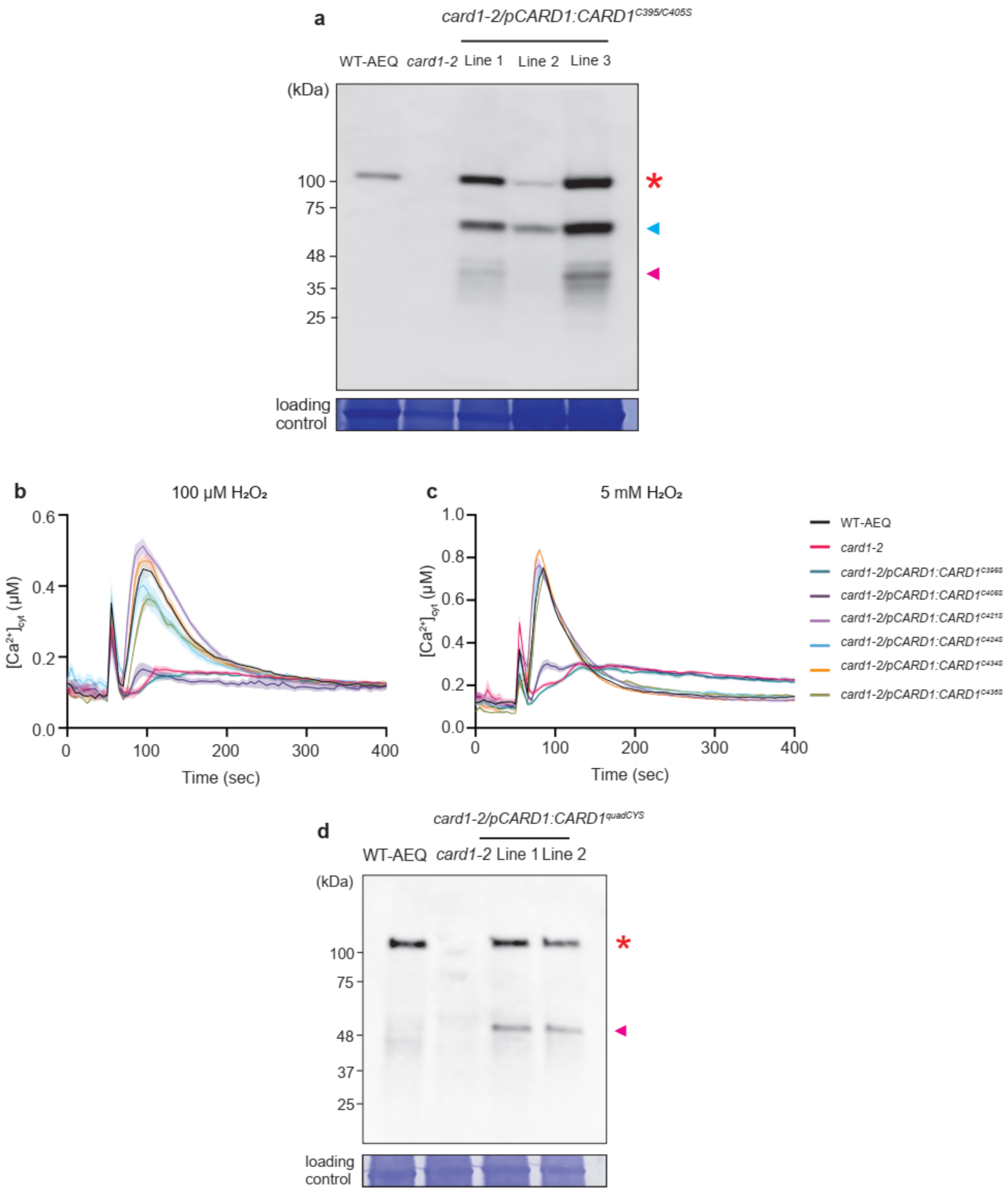

**Supplemental Figure 4. $H_2O_2$-induced $Ca^{2+}$ elevation phenotype in single cysteine amino substitution and CARD1 protein levels after mutation in the cysteine in different regions. a,** Protein levels in transgenic lines expressing CARD1 variants with Cys395 and Cys405 mutations. The red asterisk indicates full-length CARD1 protein; blue and magenta arrows indicate potential

CARD1 breakdown products. **b-c,** $[Ca^{2+}]_{cyt}$ response of *Arabidopsis* seedlings WT-AEQ, *card1-2* and *card1-2* plants expressing CARD1 variants with different cysteine mutations under the control of *AtCARD1* promoter in response to (**b**) 100 μM $H_2O_2$ and (**c**) 5 mM $H_2O_2$. Stimuli were applied at 60 s. Data are mean ± SE. $n = 8$. **d,** Protein levels in transgenic lines expressing CARD1 variants with mutations at Cys421, Cys424, Cys434, and Cys436 (quadCYS). The red asterisk indicates full-length CARD1 protein; the magenta arrow indicates a potential CARD1-derived fragment.

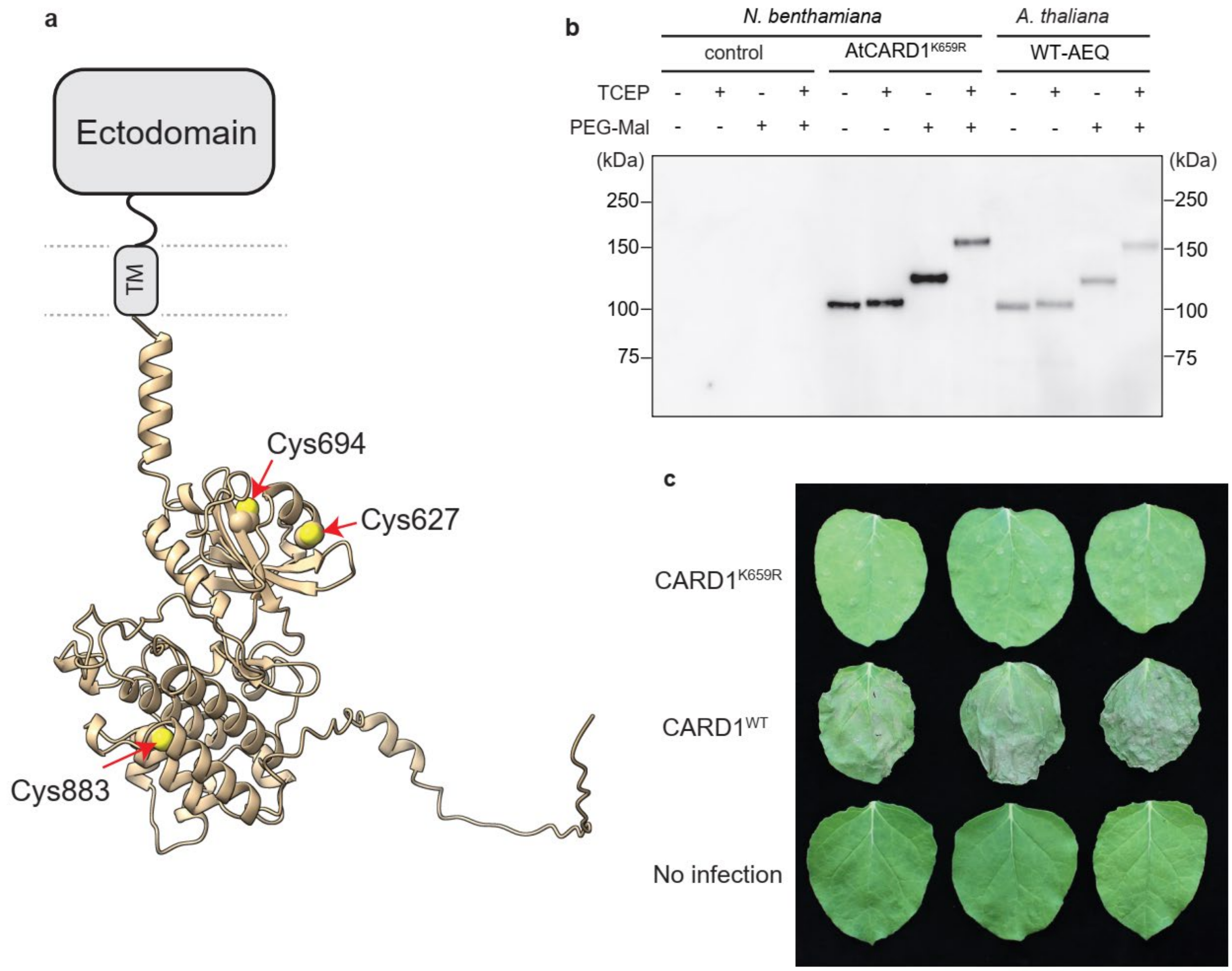

**Supplemental Figure 5. Validation of PEG-Mal labeling of cysteines in *Nicotiana benthamiana*-expressed CARD1 protein. a,** Alphafold model of CARD1 kinase domain. Cys627, Cys694, and Cys883 positions are indicated. **b,** Migration patterns of CARD1 protein during each step of PEG-Mal labeling. Protein extracts from untreated WT-AEQ or *N. benthamiana* leaves expressing a kinase-dead CARD1 variant (CARD1$^{K659R}$) were analyzed. The blot was probed with anti-CARD1 antibody. **c,** Full-length CARD1 protein induces a cell death phenotype in *N. benthamiana*, which is attenuated by mutation of Lys659.

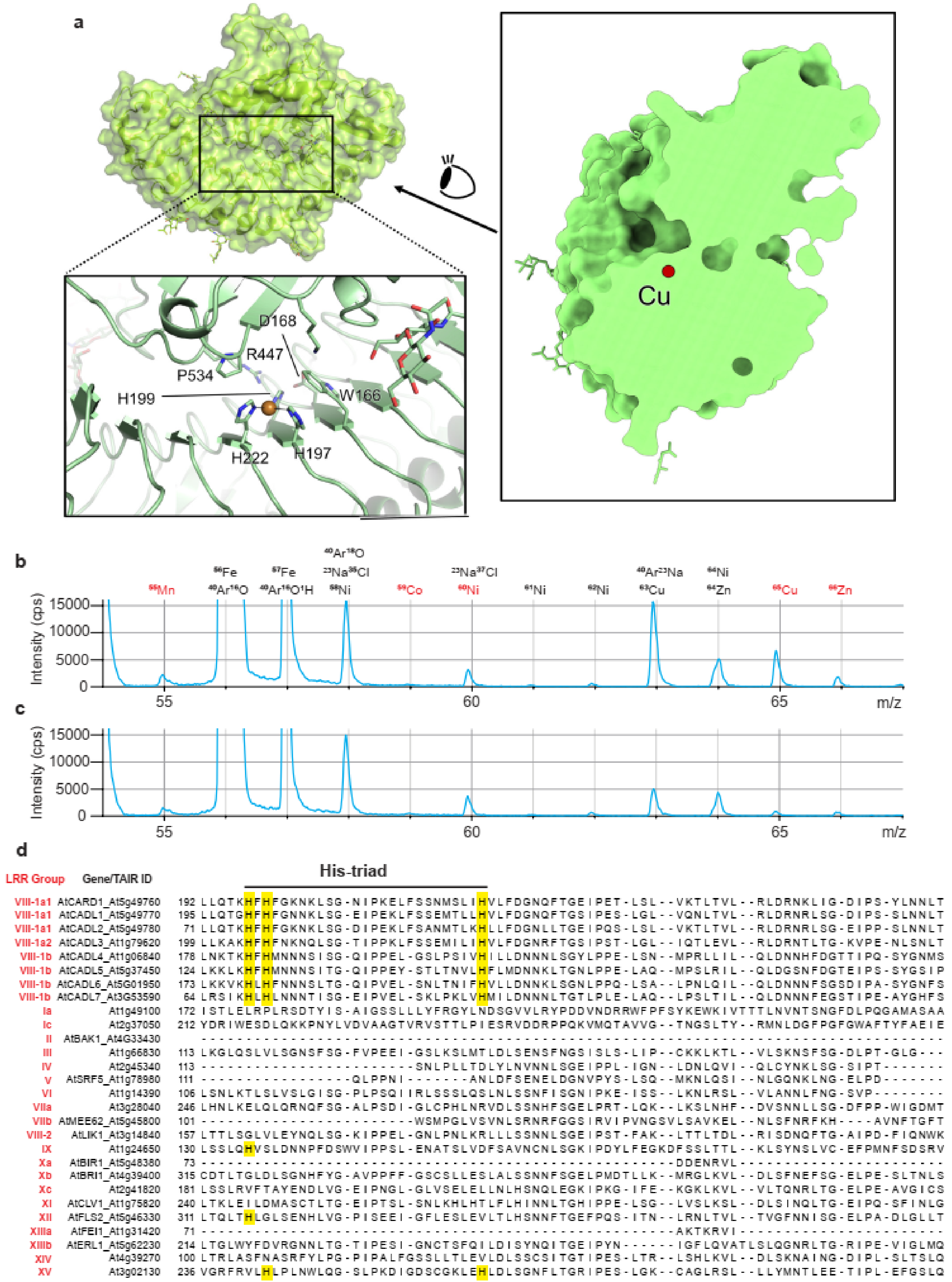
a
D168
R447
P534
H199
W166
H222
H197
Cu
b
55Mn
56Fe
40Ar16O
57Fe
40Ar16O1H
40Ar18O
23Na35Cl
58Ni
59Co
23Na37Cl
60Ni
61Ni
62Ni
40Ar23Na
63Cu
64Ni
64Zn
65Cu
66Zn
Intensity (cps)
15000
10000
5000
0
55
60
65
m/z
c
Intensity (cps)
15000
10000
5000
0
55
60
65
m/z
d
His-triad
LRR Group
Gene/TAIR ID
VIII-1a1 AtCARD1_At5g49760 192 LLQTKHFHFGKNKLSG-NIPKELFSSNMSLIHVLFDGNQFTGEIPET-LSL--VKTLTVL--RLDRNKLIG-DIPS-YLNNLT
VIII-1a1 AtCADL1_At5g49770 195 LLQTGHFHFGNNKLSG-EIPEKLFSSEMTLLHVLFDGNQFTGSIPES-LGL--VQNLTVL--RLDRNRLSG-DIPS-SLNNLT
VIII-1a1 AtCADL2_At5g49780 71 LLQTKHFHFGKNKLSG-DIPEKLFSANMTLKHLLFDGNLLTGEIPQS-LSL--VKTLTVL--RLDRNRLSG-EIPP-SLNNLT
VIII-1a2 AtCADL3_At1g79620 199 LLKAKHFHFNKNQLSG-TIPPKLFSSEMILIHVLFDGNRFTGSIPST-LGL--IQTLEVL--RLDRNTLTG-KVPE-NLSNLT
VIII-1b AtCADL4_At1g06840 178 LNKTKHFHMNNNSISG-QIPPEL-GSLPSIVHILLDNNNLSGYLPPE-LSN--MPRLLIL--QLDNNHFDGTTIPQ-SYGNMS
VIII-1b AtCADL5_At5g37450 124 LKKLKHFHMNNNSITG-QIPPEY-STLTNVLHFLMDNNKLTGNLPPE-LAQ--MPSLRIL--QLDGSNFDGTEIPS-SYGSIP
VIII-1b AtCADL6_At5G01950 173 LKKVKHLHFNNNSLTG-QIPVEL-SNLTNIFHVLLDNNKLSGNLPPQ-LSA--LPNLQIL--QLDNNNFSGSDIPA-SYGNFS
VIII-1b AtCADL7_At3G53590 64 LRSIKHLHLNNNTISG-EIPVEL-SKLPKLVHMILDNNNLTGTLPLE-LAQ--LPSLTIL--QLDNNNFEGSTIPE-AYGHFS
Ia At1g49100 172 ISTLELRPLRSDTYIS-AIGSSLLLYFRGYLNDSGVVLRYPDDVNDRRWFPFSYKEWKIVTTTLNVNTSNGFDLPQGAMASAA
Ic At2g37050 212 YDRIWESDLQKKPNYLVDVAAGTVRVSTTLPIESRVDDRPPQKVMQTAVVG--TNGSLTY--RMNLDGFPGFGWAFTYFAEIE
II AtBAK1_At4G33430
III At1g66830 113 LKGLQSLVLSGNSFSG-FVPEEI-GSLKSLMTLDLSENSFNGSISLS-LIP--CKKLKTL--VLSKNSFSG-DLPT-GLG---
IV At2g45340 113 -----------------------SNLPLLTDLYLNVNNLSGEIPPL-IGN--LDNLQVI--QLCYNKLSG-SIPT-------
V AtSRF5_At1g78980 111 ------------------QLPPNI-------ANLDFSENELDGNVPYS-LSQ--MKNLQSI--NLGQNKLNG-ELPD-------
VI At1g14390 106 LSNLKTLSLVSLGISG-PLPSQIIRLSSSLQSLNLSSNFISGNIPKE-ISS--LKNLRSL--VLANNLFNG-SVP--------
VIIa At3g28040 246 LHNLKELQLQRNQFSG-ALPSDI-GLCPHLNRVDLSSNHFSGELPRT-LQK--LKSLNHF--DVSNNLLSG-DFPP-WIGDMT
VIIb AtMEE62_At5g45800 101 -----------------------WSMPGLVSVNLSRNRFGGSIRVIPVNGSVLSAVKEL--NLSFNRFKH----AVNFTGFT
VIII-2 AtLIK1_At3g14840 157 LTTLSGLVLEYNQLSG-KIPPEL-GNLPNLKRLLLSSNNLSGEIPST-FAK--LTTLTDL--RISDNQFTG-AIPD-FIQNWK
IX At1g24650 130 LSSLQHVSLDNNPFDSWVIPPSL-ENATSLVDFSAVNCNLSGKIPDYLFEGKDFSSLTTL--KLSYNSLVC-EFPMNFSDSRV
Xa AtBIR1_At5g48380 73 ----------------------------------------------DDENRVL------------------------
Xb AtBRI1_At4g39400 315 CDTLTGLDLSGNHFYG-AVPPFF-GSCSLLESLALSSNNFSGELPMDTLLK--MRGLKVL--DLSFNEFSG-ELPE-SLTNLS
Xc At2g41820 181 LSSLRVFTAYENDLVG-EIPNGL-GLVSELELLNLHSNQLEGKIPKG-IFE--KGKLKVL--VLTQNRLTG-ELPE-AVGICS
XI AtCLV1_At1g75820 240 LTKLEILDMASCTLTG-EIPTSL-SNLKHLHTLFLHINNLTGHIPPE-LSG--LVSLKSL--DLSINQLTG-EIPQ-SFINLG
XII AtFLS2_At5g46330 311 LTQLTHLGLSENHLVG-PISEEI-GFLESLEVLTLHSNNFTGEFPQS-ITN--LRNLTVL--TVGFNNISG-ELPA-DLGLLT
XIIIa AtFEI1_At1g31420 71 ----------------------------------------------AKTKRVI------------------------
XIIIb AtERL1_At5g62230 214 LTGLWYFDVRGNNLTG-TIPESI-GNCTSFQILDISYNQITGEIPYN------IGFLQVATLSLQGNRLTG-RIPE-VIGLMQ
XIV At4g39270 100 LTRLASFNASRFYLPG-PIPALFGSSLLTLEVLDLSSCSITGTIPES-LTR--LSHLKVL--DLSKNAING-DIPL-SLTSLQ
XV At3g02130 236 VGRFRVLHLPLNWLQG-SLPKDIGDSCGKLEHLDLSGNFLTGRIPES-LGK--CAGLRSL--LLYMNTLEE-TIPL-EFGSLQ

**Supplemental Figure 6. EctoCARD1 harbors a copper ion in an exposed pocket of the protein by the conserved histidine triad (His-triad). a,** Surface-filled model of the EctoCARD1 protein. The black rectangle highlights the magnified region containing the copper ion. Right; a side view of EctoCARD1 protein and the position of $Cu^{+}$ ion relative to the surface cleft. **b-c**, Raw ICP-MS traces showing metal content in (**b**) the EctoCARD1 sample and (**c**) the buffer control. **d**, The histidine triad (His-triad) is conserved within the LRR-RLK subfamily VIII-1 but not in other LRR-RLK groups. Partial sequence alignment of CARD1 and its homologues (CADLs; subgroups VIII-1a and VIII-1b) with representative members from other LRR-RLK subfamilies. Sequence alignment was performed using MUSCLE with default parameters. Conserved histidine residues are highlighted in yellow. LRR subfamily classification is indicated in red next to each representative.

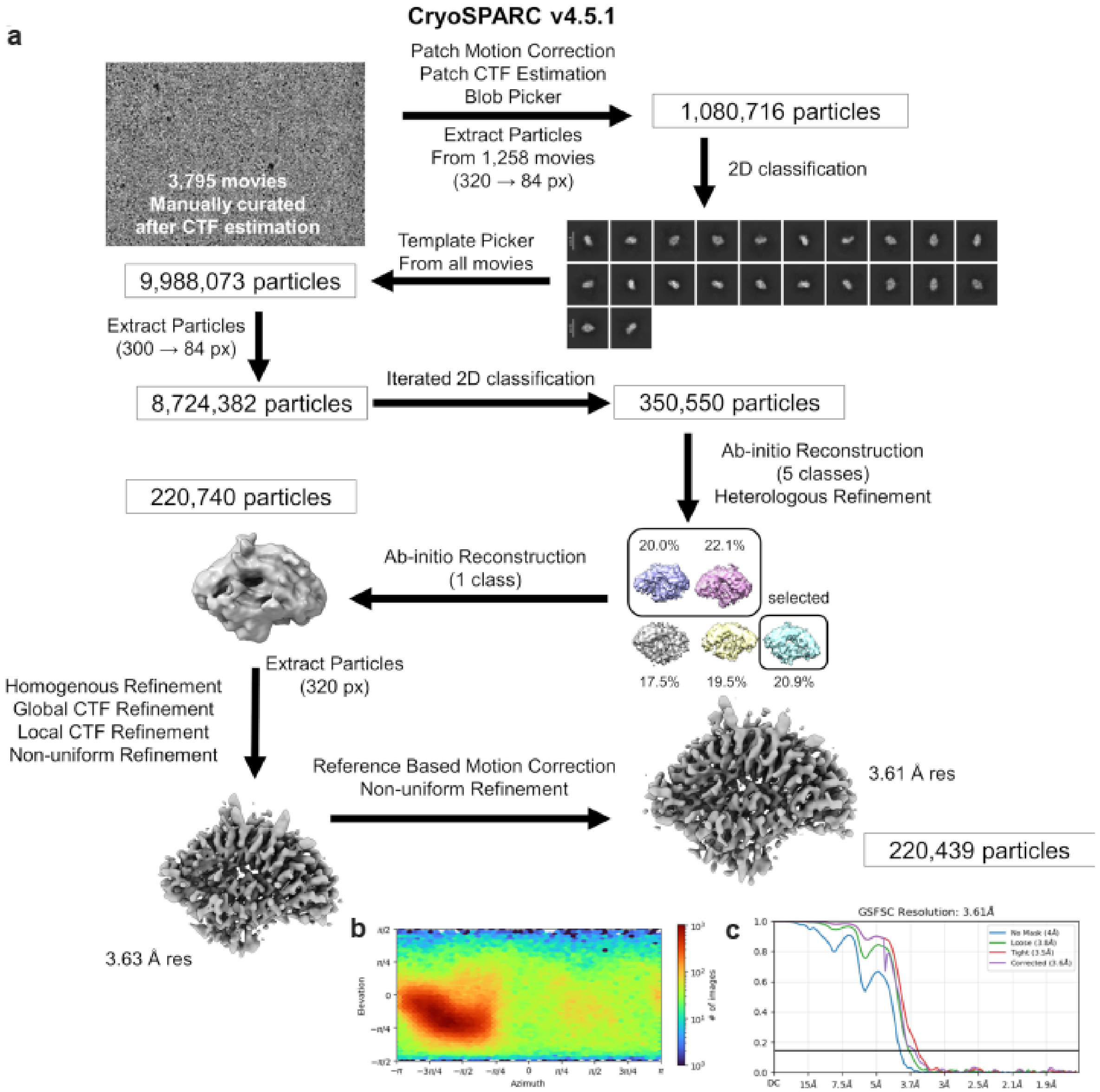

**Supplemental Figure 7. Cryo-EM data processing for EctoCARD1$^{\mathrm{HtriA}}$. a,** Workflow of data processing**. b,** Particle orientation distribution. **c,** GSFSC curves calculated with different masks.

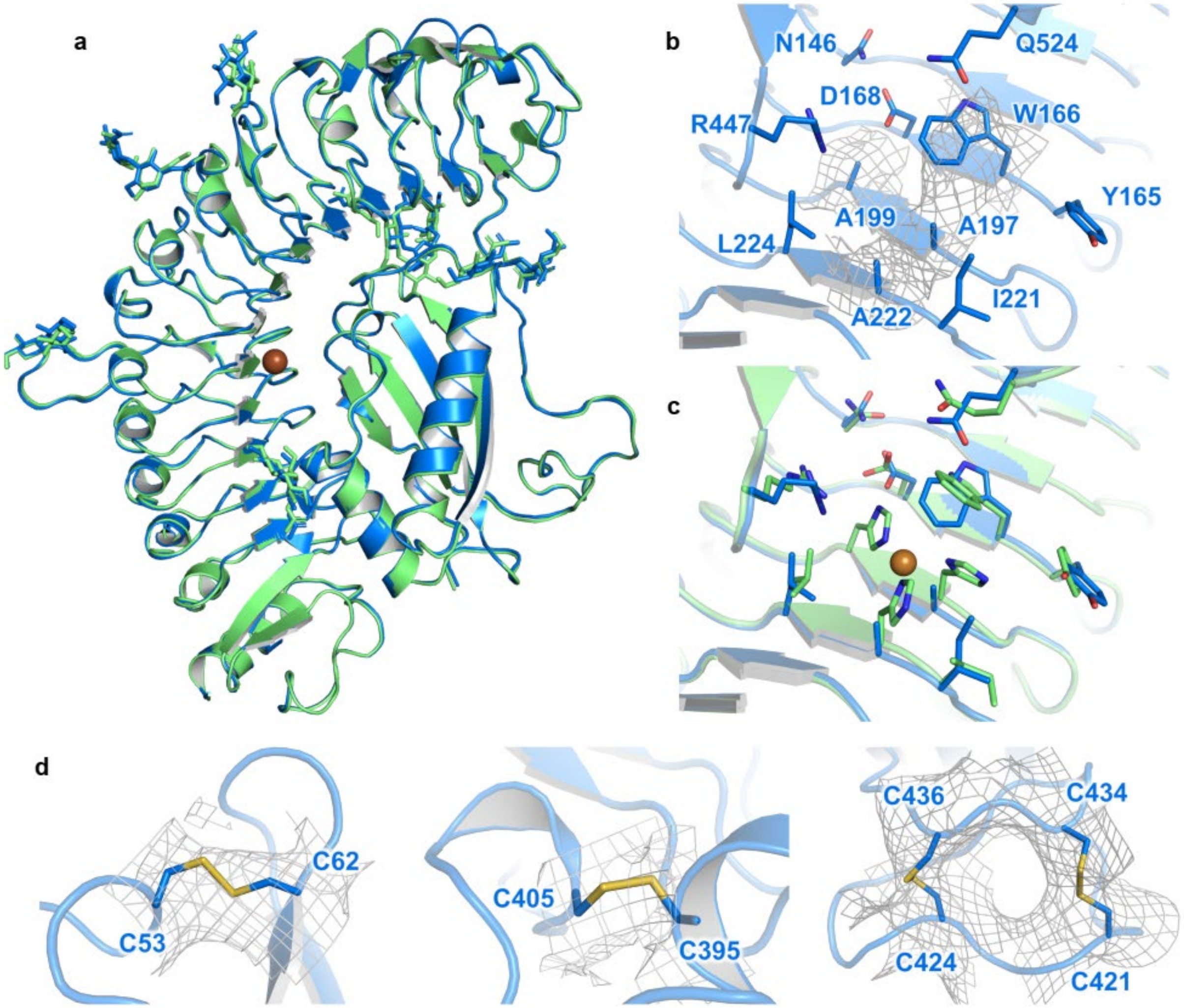

**Supplemental Figure 8. Cryo-EM structure of EctoCARD1$^{\mathbf{HtriA}}$. a,** Superimposition of EctoCARD1$^{HtriA}$ (blue) with WT EctoCARD1 (green). **b,** Mutated Cu-binding site in EctoCARD1$^{HtriA}$. The cryo-EM density maps for three alanine residues (A197, A199, and A222) and W166 are shown as gray meshes. **c,** Structural comparison of the Cu-binding site between the WT protein and EctoCARD1$^{HtriA}$. The Cu ion is shown as a brown sphere. **d,** Disulfide bonds in the EctoCARD1$^{HtriA}$ structure, with cryo-EM density maps shown by gray meshes.

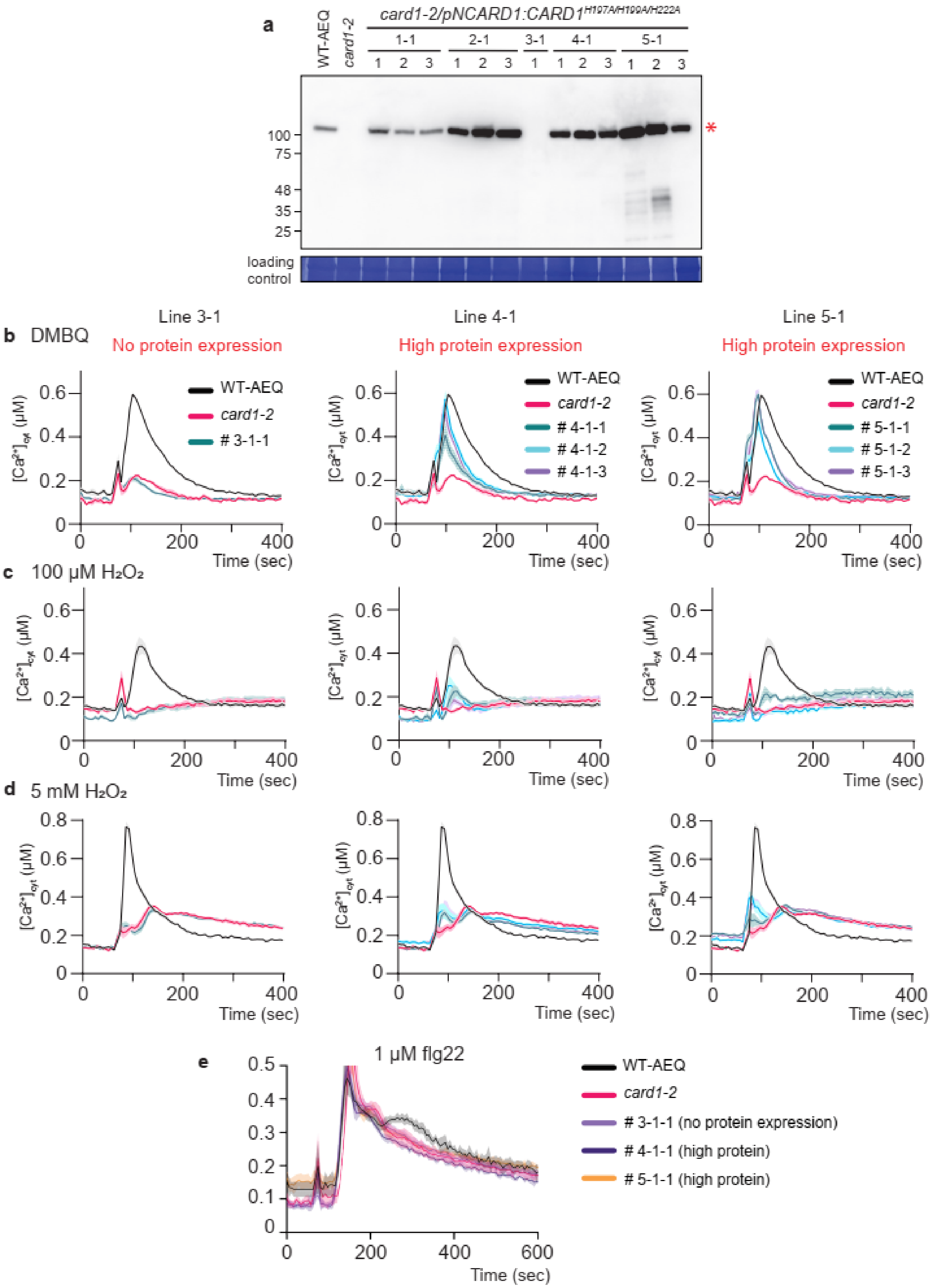

a
WT-AEQ
card1-2
card1-2/pNCARD1:CARD1H197A/H199A/H222A
1-1
2-1
3-1
4-1
5-1
1 2 3 1 2 3 1 1 2 3 1 2 3
100
75
48
35
25
*
loading control
b DMBQ
Line 3-1
No protein expression
Line 4-1
High protein expression
Line 5-1
High protein expression
WT-AEQ
card1-2
# 3-1-1
# 4-1-1
# 4-1-2
# 4-1-3
# 5-1-1
# 5-1-2
# 5-1-3
[Ca2+]cyt (µM)
Time (sec)
c 100 µM H2O2
d 5 mM H2O2
e
1 µM flg22
### 3-1-1 (no protein expression)
### 4-1-1 (high protein)
### 5-1-1 (high protein)

**Supplemental Figure 9.** Independent lines of His-triad mutants showed impaired $H_2O_2$ response. **a,** Protein levels in five independent T1 lines of CARD1 His-triad mutants (His197A/H199A/H222A), with plants grown for three generations. Three lines were selected for analysis (except line 3-1-1). Red asterisk denotes CARD1 protein. **b-d**, $[Ca^{2+}]_{cyt}$ response of *Arabidopsis* seedlings WT-AEQ, *card1-2* and T3 lines of *card1-2/CARD1^HtriA^* (from independent T1) in response to (**b**) DMBQ, (**c**) 100 µM $H_2O_2$, and (**d**) 5 mM $H_2O_2$. Stimuli were applied at 60 s. For DMBQ treatment, data are mean ± SE (*n* = 6 for WT and *card1-2*, *n* = 12 for transformant lines. For $H_2O_2$ treatments, *n* = 6 for all genotypes. **e,** $[Ca^{2+}]_{cyt}$ response of *Arabidopsis* seedlings WT-AEQ, *card1-2* and T3 lines of *card1-2/CARD1^HtriA^* (3 independent lines) in response to 1 µM flg22. Stimuli were applied at 60 s. Data are mean ± SE (*n* = 6).

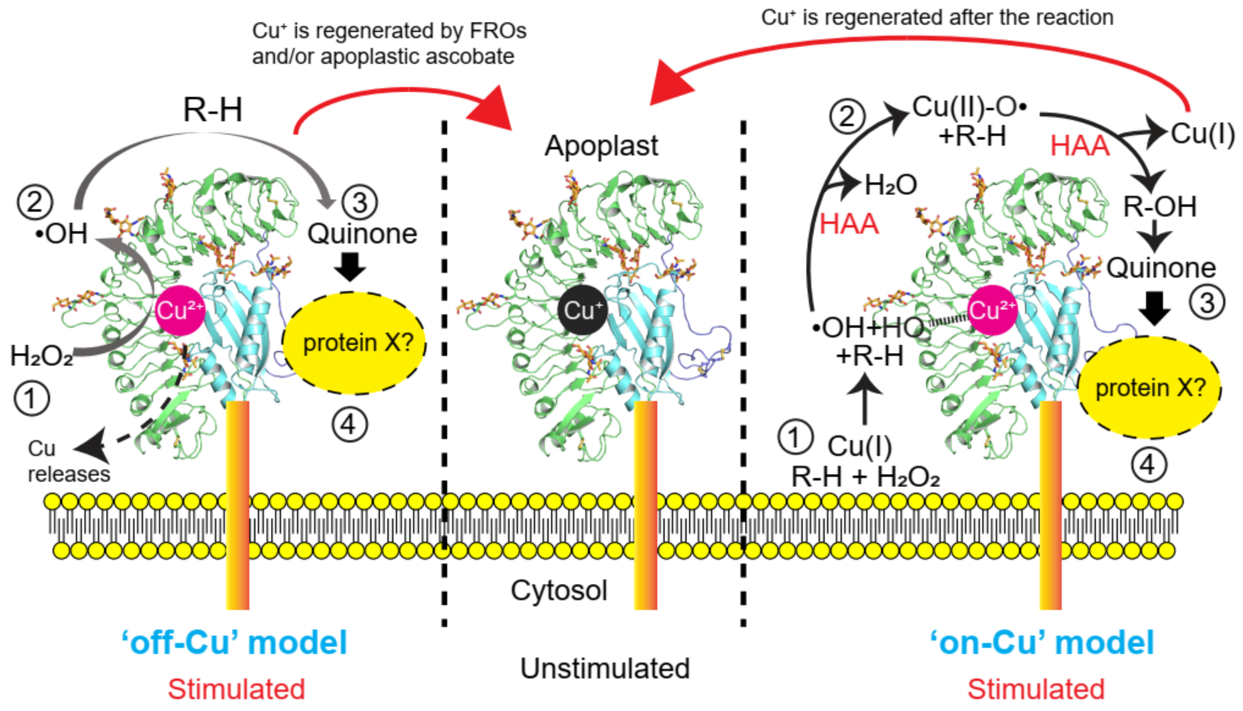

**Supplemental Figure 10. Proposed working model for CARD1 function.** In the unstimulated state, CARD1 contains $Cu^{+}$ coordinated by the His-triad. Upon stimulation with $H_2O_2$, two possible models are proposed. Left, the 'off-Cu' model. Electron transfer occurs from $Cu^{+}$ to $H_2O_2$, oxidizing $Cu^{+}$ to $Cu^{2+}$, which may subsequently dissociate from the protein ①. Concurrently, hydroxyl radicals ($^{\bullet}OH$) are generated through a Fenton reaction between $H_2O_2$ and $Cu^{+}$ in the apoplastic space ②. Due to their high reactivity, $^{\bullet}OH$ radicals can oxidize various substrates (R-H), including lignin, leading to the formation of quinones ③. These quinones may be recognized by CARD1 through a distinct region and mechanism, separate from $H_2O_2$ sensing. Additionally, other proteins or co-receptors may facilitate quinone binding *in vivo* ④. Right, the 'on-Cu' model. In this model, $Cu^{+}$ reacts with substrates – $H_2O_2$ and unknown substrate (R-H)①, This leads to the formation of a reactive copper-oxyl ($^{\bullet}O\text{-}Cu^{2+}$) intermediate *via* the first hydrogen atom abstraction (HAA) ②. After a subsequent HAA from a substrate (R-H) that is catalyzed by the $^{\bullet}O\text{-}Cu^{2+}$ intermediate, hydroxylated product (R-OH) could be generated. Subsequent reactions may generate quinones ③, which are perceived by CARD1 through the same mechanism proposed in the 'off-Cu' model ④. Note that specific substrate will be required for the 'on-Cu' model, similar to how LPMO enzyme works. This is in contrast to the 'off-Cu' model where OH radicals can attack a broader substrate. One substrate for 'on-Cu' model is hydroquinone which could be oxidized to quinone. $Cu^{+}$ is regenerated either by the ascorbate present in the apoplast or *via* ferric reductase oxidases (FROs) in plants.

**Supplementary Table 1. Cryo-EM data collection and refinement statistics.**

| | **Wild type (EMD-64920) (PDB 9VBD)** | **HtriA (EMD-64934) (PDB 9VBV)** |
|---|---|---|
| | **Data Collection** | |
| **Magnification** | 60,000 | 60,000 |
| **Voltage (kV)** | 200 | 200 |
| **Electron exposure ($e^{-}$/Å$^2$)** | 40 | 40 |
| **Defocus range (µm)** | −0.7 to −2.2 | −0.7 to −2.2 |
| **Pixel size (Å)** | 0.83 | 0.83 |
| **Symmetry imposed** | *C*1 | *C*1 |
| **Initial particle images (no.)** | 1,104,287 | 8,724,382 |
| **Final particle images (no.)** | 130,138 | 220,439 |
| **Map resolution (Å)** | 3.26 | 3.61 |
| **FSC threshold** | 0.143 | 0.143 |
| **Map resolution range (Å)** | 3.00-5.47 | 3.06-4.92 |
| | **Refinement** | |
| **Initial model used** | AlphaFold database model | Wild type structure |
| **Model resolution (Å)** | 3.56 | 3.89 |
| **FSC threshold** | 0.5 | 0.5 |
| **Map sharpening *B* factor (Å$^2$)** | 147.2 | 162.8 |
| **Model composition** | | |
| **Non-hydrogen atoms** | 4243 | 4241 |
| **Protein residues** | 521 | 521 |
| ***B* factors (Å)** | | |
| **Protein** | 62.8 | 35.9 |
| **Ligand** | 86.3 | 64.8 |
| **R.m.s. deviations** | | |
| **Bond lengths (Å)** | 0.003 | 0.004 |
| **Bond angles (°)** | 0.599 | 0.662 |
| **Ramachandran plot** | | |
| **Favored (%)** | 95.18 | 94.41 |
| **Allowed (%)** | 4.82 | 5.59 |
| **Disallowed (%)** | 0.00 | 0.00 |

**Supplementary Table 2. Comparison of SEA domains with Z score > 5 in Dali.**

| No | PDBID-Chain | Z score | rmsd (Å) | lali | nres | %id | PDB Description |
|---|---|---|---|---|---|---|---|
| **1** | 2e7v-A | 8.1 | 2.5 | 84 | 115 | 11 | MOLECULE: TRANSMEMBRANE PROTEASE; |
| **2** | 6ffy-A | 7.8 | 2.7 | 90 | 918 | 13 | MOLECULE: VPS10 DOMAIN-CONTAINING RECEPTOR SORCS2; |
| **3** | 8vm1-A | 7.8 | 3.2 | 95 | 185 | 11 | MOLECULE: MESOTHELIN, CLEAVED FORM,MUCIN-16; |
| **4** | 6v55-A | 7.7 | 3 | 90 | 746 | 9 | MOLECULE: ADHESION G-PROTEIN COUPLED RECEPTOR G6; |
| **5** | 6eet-A | 7 | 3 | 90 | 443 | 8 | MOLECULE: PROTOCADHERIN-15; |
| **6** | 7wwu-B | 7 | 3 | 87 | 248 | 8 | MOLECULE: CSY1; |
| **7** | 5ijn-F | 7 | 2.9 | 84 | 335 | 14 | MOLECULE: NUCLEAR PORE COMPLEX PROTEIN NUP155; |
| **8** | 8tdl-A | 6.4 | 3.4 | 83 | 364 | 10 | MOLECULE: MECHANOSENSITIVE ION CHANNEL PROTEIN 10; |
| **9** | 5ehk-A | 6.3 | 3.2 | 81 | 1036 | 2 | MOLECULE: LANTIBIOTIC DEHYDRATASE; |
| **10** | 6vqv-D | 6.1 | 3.5 | 87 | 306 | 14 | MOLECULE: ACRF9; |
| **11** | 6zyd-A | 6 | 3.1 | 80 | 323 | 10 | MOLECULE: LOW CONDUCTANCE MECHANOSENSITIVE CHANNEL YNAI,LOW |
| **12** | 6exx-A | 5.9 | 2.2 | 66 | 79 | 9 | MOLECULE: PROTEIN PES4; |
| **13** | 8tdj-A | 5.9 | 3.3 | 82 | 467 | 10 | MOLECULE: MECHANOSENSITIVE ION CHANNEL PROTEIN 10; |
| **14** | 5cio-A | 5.9 | 3.1 | 88 | 770 | 7 | MOLECULE: PYRROLOQUINOLINE QUINONE BIOSYNTHESIS PROTEIN PQQ |
| **15** | 3mcs-B | 5.8 | 4.1 | 83 | 218 | 7 | MOLECULE: PUTATIVE MONOOXYGENASE; |
| **16** | 7u5d-A | 5.8 | 3.1 | 91 | 630 | 7 | MOLECULE: CRRNA; |
| **17** | 6vxm-A | 5.8 | 3.2 | 84 | 277 | 5 | MOLECULE: MECHANOSENSITIVE ION CHANNEL PROTEIN 1, MITOCHOND |
| **18** | 1yz7-A | 5.8 | 3.2 | 83 | 176 | 7 | MOLECULE: PROBABLE TRANSLATION INITIATION FACTOR 2 ALPHA |
| **19** | 8ecx-C | 5.8 | 3.7 | 82 | 102 | 9 | MOLECULE: ANTIBIOTIC BIOSYNTHESIS MONOOXYGENASE; |
| **20** | 1r3n-A | 5.8 | 3.8 | 83 | 438 | 8 | MOLECULE: BETA-ALANINE SYNTHASE; |
| **21** | 4wd9-A | 5.7 | 3.5 | 79 | 965 | 11 | MOLECULE: NISIN BIOSYNTHESIS PROTEIN NISB; |
| **22** | 8d97-A | 5.7 | 3.4 | 86 | 1600 | 6 | MOLECULE: RAMP SUPERFAMILY PROTEIN; |
| **23** | 6fjw-A | 5.7 | 3.2 | 79 | 239 | 4 | MOLECULE: CAS6 PROTEIN; |
| **24** | 6wub-f | 5.7 | 2.9 | 80 | 97 | 9 | MOLECULE: 16S RRNA; |
| **25** | 7db7-B | 5.7 | 3.2 | 84 | 836 | 8 | MOLECULE: PHENYLALANINE--TRNA LIGASE ALPHA SUBUNIT; |
| **26** | 2raq-B | 5.7 | 2.5 | 67 | 94 | 7 | MOLECULE: CONSERVED PROTEIN MTH889; |
| **27** | 8vrc-A | 5.6 | 2.4 | 70 | 100 | 9 | MOLECULE: LA MOTIF RNA-BINDING DOMAIN PROTEIN; |

| No | PDBID-Chain | Z score | rmsd (Å) | lali | nres | %id | PDB Description |
|---|---|---|---|---|---|---|---|
| **28** | 6slf-A | 5.6 | 3.9 | 84 | 398 | 8 | MOLECULE: N-ALPHA-ACYL-GLUTAMINE AMINOACYLASE; |
| **29** | 1q8b-A | 5.6 | 3.7 | 81 | 93 | 9 | MOLECULE: PROTEIN YJCS; |
| **30** | 1k8w-A | 5.6 | 2.6 | 75 | 303 | 5 | MOLECULE: 5'-R(*GP*GP*CP*AP*AP*CP*GP*GP*UP*(FHU) |
| **31** | 4v1a-k | 5.6 | 2.8 | 75 | 131 | 4 | MOLECULE: MITORIBOSOMAL PROTEIN ML37, MRPL37; |
| **32** | 2od4-A | 5.6 | 3 | 76 | 101 | 4 | MOLECULE: HYPOTHETICAL PROTEIN; |
| **33** | 6urt-A | 5.6 | 3 | 81 | 331 | 9 | MOLECULE: LOW CONDUCTANCE MECHANOSENSITIVE CHANNEL YNAI; |
| **34** | 6fxd-A | 5.5 | 3.6 | 77 | 128 | 6 | MOLECULE: MUPZ; |
| **35** | 7r9g-A | 5.5 | 3.6 | 84 | 386 | 7 | MOLECULE: TRNA PSEUDOURIDINE SYNTHASE 1; |
| **36** | 7yzi-A | 5.5 | 3.4 | 84 | 378 | 6 | MOLECULE: ADENYLATE CYCLASE; |
| **37** | 1j27-A | 5.4 | 3.1 | 82 | 98 | 7 | MOLECULE: HYPOTHETICAL PROTEIN TT1725; |
| **38** | 6wi5-A | 5.4 | 3.8 | 78 | 90 | 10 | MOLECULE: DE NOVO DESIGNED PROTEIN FOLDIT4; |
| **39** | 4x4r-A | 5.4 | 2.8 | 86 | 443 | 9 | MOLECULE: CCA-ADDING ENZYME; |
| **40** | 5mgu-A | 5.3 | 3.3 | 80 | 408 | 9 | MOLECULE: PHENYLALANINE--TRNA LIGASE, MITOCHONDRIAL; |
| **41** | 6w6v-E | 5.3 | 3 | 80 | 169 | 5 | MOLECULE: RNA COMPONENT OF RNASE MRP NME1; |
| **42** | 7c50-A | 5.3 | 3.4 | 84 | 192 | 11 | MOLECULE: SIMPL DOMAIN-CONTAINING PROTEIN; |
| **43** | 2hiy-D | 5.3 | 2.9 | 67 | 183 | 6 | MOLECULE: HYPOTHETICAL PROTEIN; |
| **44** | 9iad-A | 5.3 | 2.6 | 67 | 685 | 9 | MOLECULE: PROTEIN ARGONAUTE; |
| **45** | 9go3-A | 5.3 | 3 | 81 | 253 | 11 | MOLECULE: MECHANOSENSITIVE CHANNEL PROTEIN; |
| **46** | 7bio-B | 5.2 | 3.7 | 74 | 111 | 9 | MOLECULE: MONOOXYGENASE/PUTATIVE ANTHRONOXYGENASE; |
| **47** | 6wu0-B | 5.2 | 3.5 | 80 | 854 | 8 | MOLECULE: HOPANOID BIOSYNTHESIS ASSOCIATED RND TRANSPORTER |
| **48** | 6nuk-A | 5.2 | 3.8 | 76 | 101 | 5 | MOLECULE: FERREDOG-DIESEL; |
| **49** | 6v3f-A | 5.2 | 4 | 92 | 1234 | 8 | MOLECULE: NPC1-LIKE INTRACELLULAR CHOLESTEROL TRANSPORTER 1 |
| **50** | 8h2h-D | 5.2 | 3.1 | 82 | 599 | 9 | MOLECULE: LTRB; |
| **51** | 6vej-A | 5.1 | 3.3 | 79 | 1022 | 9 | MOLECULE: PROBABLE RESISTANCE-NODULATION-CELL DIVISION (RND |
| **52** | 4p6q-A | 5.1 | 2.4 | 69 | 286 | 12 | MOLECULE: MSX2-INTERACTING PROTEIN; |
| **53** | 7xjz-A | 5.1 | 3.2 | 83 | 983 | 10 | MOLECULE: GLYCINE--TRNA LIGASE; |
| **54** | 2yy3-A | 5.1 | 3 | 77 | 91 | 8 | MOLECULE: ELONGATION FACTOR 1-BETA; |
| **55** | 7n7s-B | 5.1 | 3 | 89 | 441 | 10 | MOLECULE: HYDROXYMETHYLGLUTARYL-COA REDUCTASE; |
| **56** | 3gz7-A | 5.1 | 3.7 | 82 | 99 | 9 | MOLECULE: PUTATIVE ANTIBIOTIC BIOSYNTHESIS MONOOXYGENASE; |

| No | PDBID-Chain | Z score | rmsd (Å) | lali | nres | %id | PDB Description |
|---|---|---|---|---|---|---|---|
| **57** | 4djb-A | 5.1 | 4.4 | 86 | 118 | 10 | MOLECULE: E4-ORF3; |
| **58** | 1y7p-B | 5.1 | 2.9 | 69 | 217 | 9 | MOLECULE: HYPOTHETICAL PROTEIN AF1403; |
| **59** | 7mp7-A | 5.1 | 3.2 | 77 | 87 | 6 | MOLECULE: SB3; |
| **60** | 2x3g-A | 5 | 3 | 76 | 116 | 11 | MOLECULE: SIRV1 HYPOTHETICAL PROTEIN ORF119; |
| **61** | 3bm7-A | 5 | 4.2 | 84 | 106 | 7 | MOLECULE: PROTEIN OF UNKNOWN FUNCTION WITH FERREDOXIN-LIKE |
| **62** | 7szy-A | 5 | 2.9 | 84 | 174 | 10 | MOLECULE: NS1 PROTEIN; |
| **63** | 6rie-A | 5 | 3.4 | 75 | 1285 | 5 | MOLECULE: DNA-DEPENDENT RNA POLYMERASE SUBUNIT RPO147; |
| **64** | 9dp6-A | 5 | 3.2 | 79 | 737 | 10 | MOLECULE: MEROMYCOLATE EXTENSION ACYL CARRIER PROTEIN; |
| **65** | 3pfe-A | 5 | 3.6 | 79 | 471 | 16 | MOLECULE: SUCCINYL-DIAMINOPIMELATE DESUCCINYLASE; |
| **66** | 6f7s-C | 5 | 2.6 | 71 | 319 | 3 | MOLECULE: SERRATE RNA EFFECTOR MOLECULE HOMOLOG; |
| **67** | 5yhf-A | 5 | 3.3 | 78 | 734 | 4 | MOLECULE: PROTEIN TRANSLOCASE SUBUNIT SECDF; |
| **68** | 8wg3-A | 5 | 2.8 | 73 | 548 | 5 | MOLECULE: CSC1-LIKE PROTEIN 2,GREEN FLUORESCENT PROTEIN; |
| **69** | 3al0-B | 5 | 3.4 | 91 | 482 | 10 | MOLECULE: GLUTAMYL-TRNA(GLN) AMIDOTRANSFERASE SUBUNIT A; |
| **70** | 2f7v-A | 5 | 4.1 | 81 | 360 | 7 | MOLECULE: AECTYLCITRULLINE DEACETYLASE; |
| **71** | 2pd1-A | 5 | 3.5 | 81 | 103 | 2 | MOLECULE: HYPOTHETICAL PROTEIN; |
| **72** | 6u9d-L | 5 | 3.1 | 75 | 255 | 5 | MOLECULE: ACETOLACTATE SYNTHASE CATALYTIC SUBUNIT, MITOCHON |
| **73** | 5mz2-D | 5 | 3 | 86 | 482 | 3 | MOLECULE: RUBISCO LARGE SUBUNIT; |
| **74** | 5euf-B | 5 | 3.6 | 83 | 416 | 5 | MOLECULE: PROTEASE; |
| **75** | 2dgx-A | 5 | 2.7 | 71 | 96 | 11 | MOLECULE: KIAA0430 PROTEIN; |
| **76** | 7uoi-A | 5 | 3.8 | 75 | 383 | 7 | MOLECULE: SUCCINYL-DIAMINOPIMELATE DESUCCINYLASE; |
| **77** | 8d8k-J | 5 | 2.9 | 79 | 144 | 8 | MOLECULE: PROBABLE S-ADENOSYL-L-METHIONINE-DEPENDENT RNA |

**Supplementary Table 3. Comparison of SEA domains with top 15 entries in Foldseek.**

| Rank | Target | Description | Scientific Name | Prob | Seq. Id. | E-Value | Score | Query Pos. | Target Pos. |
|---|---|---|---|---|---|---|---|---|---|
| **1** | 7sa9-assembly2_B | Human MUC16 SEA5 Domain | Homo sapiens | 0.97 | 11.9 | 1.71E-02 | 76 | 3-102 (103) | 7-107 (120) |
| **2** | 5mgu-assembly1_A | Kinetic and Structural Changes in HsmtPheRS, Induced by Pathogenic Mutations in Human FARS2 | Homo sapiens | 0.94 | 6.8 | 3.95E-02 | 71 | 3-103 (103) | 322-407 (407) |
| **3** | 5mgh-assembly1_A | Crystal structure of pathogenic mutants of human mitochodnrial PheRS | Homo sapiens | 0.93 | 7.8 | 3.05E-02 | 70 | 3-103 (103) | 320-405 (405) |
| **4** | 8p8x-assembly1_A | Crystal structure of a pathogenic mutant variant of human mitochodnrial PheRS | Homo sapiens | 0.89 | 9.8 | 5.44E-02 | 66 | 3-103 (103) | 319-404 (404) |
| **5** | 5mgw-assembly1_A | Kinetic and Structural Changes in HsmtPheRS, Induced by Pathogenic Mutations in Human FARS2 | Homo sapiens | 0.85 | 7.4 | 4.49E-02 | 64 | 3-103 (103) | 321-406 (406) |
| **6** | 6v55-assembly1_A | Full extracellular region of zebrafish Gpr126/Adgrg6 | Danio rerio | 0.8 | 13.2 | 2.52E-02 | 61 | 3-103 (103) | 330-418 (746) |
| **7** | 3tup-assembly1_A | Crystal structure of human mitochondrial PheRS complexed with tRNAPhe in the active open state | Homo sapiens | 0.72 | 11.6 | 1.10E-01 | 58 | 3-87 (103) | 319-394 (404) |
| **8** | 3mgj-assembly1_A | Crystal structure of the Saccharop_dh_N domain of MJ1480 protein from Methanococcus jannaschii. Northeast Structural Genomics Consortium Target MjR83a. | Methanocaldococcus jannaschii | 0.6 | 10.2 | 2.54E-01 | 54 | 1-103 (103) | 1-84 (97) |
| **9** | 3rg6-assembly1_B | Crystal structure of a chaperone-bound assembly intermediate of form I Rubisco | Synechococcus elongatus PCC 6301 | 0.57 | 15.1 | 1.42E-01 | 53 | 2-83 (103) | 16-96 (438) |

| Rank | Target | Description | Scientific Name | Prob | Seq. Id. | E-Value | Score | Query Pos. | Target Pos. |
|---|---|---|---|---|---|---|---|---|---|
| **10** | 8io2-assembly1_E | The Rubisco assembly intermediate of Arabidopsis thaliana Rubisco accumulation factor 1 (AtRaf1) and Rubisco large subunit (RbcL) | Synechococcus elongatus PCC 6301 | 0.51 | 15 | 2.54E-01 | 51 | 2-90 (103) | 17-99 (415) |
| **11** | 5mgv-assembly1_A | Kinetic and Structural Changes in HsmtPheRS, Induced by Pathogenic Mutations in Human FARS2 | Homo sapiens | 0.47 | 9.8 | 3.98E-01 | 50 | 3-82 (103) | 320-390 (405) |
| **12** | 2e7v-assembly1_A | Crystal structure of SEA domain of transmembrane protease from Mus musculus | Mus musculus | 0.44 | 9.2 | 1.04E+00 | 49 | 2-89 (103) | 6-96 (115) |
| **13** | 3obi-assembly1_A | Crystal structure of a formyltetrahydrofolate deformylase (NP_949368) from RHODOPSEUDOMONAS PALUSTRIS CGA009 at 1.95 A resolution | Rhodopseudomonas palustris CGA009 | 0.44 | 10.7 | 7.03E-02 | 49 | 2-101 (103) | 4-82 (285) |
| **14** | 7l7f-assembly1_D | Cryo-EM structure of human ACE2 receptor bound to protein encoded by vaccine candidate BNT162b1 | Homo sapiens | 0.41 | 11.6 | 1.34E-01 | 48 | 3-87 (103) | 597-690 (711) |
| **15** | 3isz-assembly1_B | Crystal structure of mono-zinc form of succinyl-diaminopimelate desuccinylase from Haemophilus influenzae | Haemophilus influenzae Rd KW20 | 0.35 | 16 | 3.28E-01 | 46 | 2-103 (103) | 182-279 (366) |

**Supplementary Table 4. List of primers used in this study.**

| Primer name | Sequences (5’ -> 3’) |
|---|---|
| *For generation of EctoCARD1-MBP and EctoCARD1$^{HtriA}$-MBP for cryo-EM* | |
| **CARD1_F** | TAAACGTCTCTAAAAATGAGTTCAAGAACTGGAGCCTC |
| **CARD1_1-546_R** | AGAGAACTGTTTGTACAAATCAGC |
| **MBPco_F+GSA** | TACAAACAGTTCTCTGGGAGCGCTATGAAAATCGAAGAAGGTAAACTGG |
| **MBPco_R** | AATGAAACCAGAGCGCTAAGTCTGAGCGTCTTTCAGGGC |
| **CARD1_R600** | AGTTTGAAGAAGCATGTCAAGTCC |
| **CARD1_F691** | GATGGAAACCAATTCACGGGCG |
| **CARD1_HistriAla_F** | ATGCTTCTTCAAACTAAGGCTTTTGCTTTTGGAAAAAACAAGCTTTCAGGC |
| **CARD1_HistriAla_R** | GAATTGGTTTCCATCGAAGAGGACAGCTATTAAACTCATGTTTGAGCTGAAAAG |
| *For generation of N. benthamiana plants expressing mutated cysteine for PEG-Mal experiments* | |
| **CARD1_F** | TAAACGTCTCTAAAAATGAGTTCAAGAACTGGAGCCTC |
| **CARD1_R** | AATGAAACCAGAGCGTCATTGGGGCTCAAGCTTTGAAG |
| **CARD1_R_KR mut** | TTGCTATGAGTTGCCCATTAGG |
| **CARD1_F_KR mut** | GGCAACTCATAGCAATCCGAAGGGCTCAACAAGGATC |
| **CARD1_F_C421/434S** | AACCGGGCATGGAAGCGAGTCCCACGTCCCGCTGTGCGTATCCATTC |
| **CARD1_R_C421/434S** | GCTTCCATGCCCGGTTCGCATGGAGAAGAGTTTGTTGGGAGTGTAG |
| **CARD1_F_C424/436S** | GCATGGAAGCGAGTCCCACGTGCCGCTCTGCGTATCCATTCATGG |
| **CARD1_R_C424/436S** | GGACTCGCTTCCATGCCCGGTTCGGATGGAGAACAGTTTGTTGG |
| *For complementation assay of proCARD1-AtCADL1 or proCARD1-AtCADL3* | |
| **Fw CARD1 promoter** | AATCACAAGTTTGTACACGTGTTTGCCAAGGTACATC |
| **Fw_AtCADL1** | TTTTTGGGTTTGAAGATGAAGATGAGTTCAAGAATTGGATT |
| **Rv_AtCADL1** | CGCTTTACTTGTACATTAGGGTTTTGGAGTTGGGAAAAC |
| **Fw_AtCADL3** | TTTTTGGGTTTGAAGATGGTGGGTTCCAACACCGTAAC |
| **Rv_AtCADL3** | CGCTTTACTTGTACATTACTTGGGCTCAATTTTGGTCG |
| **Rv CARD1 promoter** | CTTCAAACCCAAAAAGAACCTCT |
| *For complementation assay of proCARD1-CARD1 C395S/C405S or proCARD1-CARD1 quadCYS* | |
| **Fw CARD1 promoter** | AATCACAAGTTTGTACACGTGTTTGCCAAGGTACATC |
| **C395C405_Fw** | AATCCAGTGAGTCTAGAGGCGGGAAACGGGCCGAGTTACAGTTCAGCA |
| **C395C405_Rv** | TGCTGAACTGTAACTCGGCCCGTTTCCCGCCTCTAGACTCACTGGATT |
| **quadcys_Fw** | CCAACAAACAGTTCTCCAAGCGAACCGGGCATGGAAGCGAGTCCCACGAGCCGCAGTGCGTAT |
| **quadcys_Rv** | ATACGCACTGCGGCTCGTGGGACTCGCTTCCATGCCCGGTTCGCTTGGAGAACTGTTTGTTGG |
| **AtCARD1_R ver2** | GGCCGCTTTACTTGTACATCATTGGGGCTCAAGCTTTG |
| *For cloning proCARD1-CARD1 HtriA construct* | |
| **Fw_CARD1 pro BamHI over** | ATTCGCGGTACCCGGGGATCCGTGTTTGCCAAGGTACATC |
| **Fw_CARD1_H197A** | AAACTAAGGCTTTGTATATATAATCCCAAATC |
| **RV_CARD1_H197A** | TATATACAAAGCCTTAGTTTGAAGAAGCATGTC |
| **Fw_CARD1_H199A** | TCTTGTAGTGCTTTTGGAAAAAACAAGCTTTCAG |
| **RV_CARD1_H199A** | TTTCCAAAAGCACTACAAGAAAAAAGCAAAGG |
| **Fw_CARD1_H222A** | AGTTTAATAGCTGTGTAAGCTTCATATTACATTATA |
| **RV_CARD1_H222A** | GCTTACACAGCTATTAAACTCATGTTTGAGCTG |
| **Rv_CARD1_XhoIover** | ATGTTTGAACGATCCTCGATCATTGGGGCTCAAGCTTTG |

**Supplementary Table 5. Operating conditions for ICP-TOFMS**

| **ICP** | |
|---|---|
| **Generator frequency** | 27.12 MHz |
| **RF applied power** | 1400 W |
| **Cooling gas flow rate** | 10.0 L/min |
| **Auxiliary gas flow rate** | 0.50 L/min |
| **Nebulizer gas flow rate** | 0.69 L/min |

| **TOFMS** | |
|---|---|
| **Data acquisition** | Continuous mode |
| **Detected mass range** | 6-260 m/z |
| **Removed mass range by ion blanker** | 11-45 m/z |
| **TOF integration time** | 5 s |
| **Replicates** | 5 times |
| **Internal standards** | $^{115}In$ |
| **Isotopes** | $^{55}Mn$, $^{59}Co$, $^{60}Ni$, $^{65}Cu$, $^{66}Zn$ |

**Supplementary Movie 1. MD simulation of $Cu^{+}$ binding affinity to the EctoCARD1 protein.**

**Supplementary Movie 2. MD simulation of $Cu^{2+}$ binding affinity to the EctoCARD1 protein.**

**Supplementary Data 1. EctoCARD1 sequences for cryoEM study.**

**Supplementary Data 2. List of CARD1/CADL homologs identified from Phytozome (version 12.1).**

**Supplementary Data 3. List of CARD1 homologs identified from the NCBI database.**
