## Supplemental Data 1 for "A copper-dependent, redox-based hydrogen peroxide perception in plants"

**Extended Data 1.**

**Nucleotide and amino acid sequences of EctoCARD1-MBP**

EctoCARD1-GSA linker-MBP

Nucleotide

ATGAGTTCAAGAACTGGAGCCTCTTTGCTCCTGATCTTGTTTTTCTTCCAAATTTGTTCTGTTTCGGC
ACTCACGAATGGTTTAGACGCTTCTGCTTTAAATGCTCTGAAGAGTGAATGGACCACTCCTCCTG
ATGGTTGGGAAGGCTCTGATCCTTGTGGAACCAATTGGGTTGGAATTACATGTCAAAATGACCGC
GTCGTTTCAATATCACTAGGTAACCTTGACTTGGAAGGAAAGCTTCCTGCCGATATATCGTTCTTGT
CCGAGTTGCGGATCTTGGATTTGTCATACAATCCTAAATTGTCCGGACCGCTTCCACCAAACATC
GGTAACCTTGGGAAGTTAAGGAACTTGATCCTTGTGGGATGTAGTTTCAGTGGTCAAATCCCTGA
GTCCATTGGAACTCTAAAAGAACTTATATATCTCTCCCTAAATTTAAATAAATTTAGTGGAACAATTCC
GCCTTCAATTGGCCTGTTATCAAAACTATATTGGTTTGATATAGCTGACAATCAGATTGAAGGAGAG
CTTCCAGTTTCTAATGGGACTTCTGCACCTGGACTTGACATGCTTCTTCAAACTAAGCATTTTCATT
TTGGAAAAAACAAGCTTTCAGGCAATATCCCAAAAGAACTTTTCAGCTCAAACATGAGTTTAATAC
ATGTCCTCTTCGATGGAAACCAATTCACGGGCGAAATTCCAGAGACCCTCAGTCTCGTTAAAAC
GTTGACGGTGTTACGCCTTGATAGGAACAAACTCATTGGAGATATTCCTTCATATCTTAACAATCTC
ACAAATCTTAACGAACTGTACTTGGCCAACAACAGATTTACTGGTACTCTTCCAAATTTAACCAGC
TTGACCAGTCTCTACACATTGGATGTGAGCAATAACACTTTGGATTTCTCACCCATTCCATCATGG
ATCTCTTCATTACCCTCCCTATCAACATTAAGGATGGAAGGGATCCAACTTAATGGTCCAATACCA
ATCTCATTTTTCAGCCCTCCTCAGTTGCAGACTGTTATCTTAAAGCGAAATAGTATAGTGGAAAGCT
TAGACTTTGGTACAGACGTTAGCAGCCAGTTGGAGTTTGTGGATTTACAATACAATGAAATAACTGA
TTATAAACCATCAGCTAATAAAGTCCTCCAAGTAATATTGGCAAATAATCCAGTGTGTCTAGAGGCG
GGAAACGGGCCGAGTTACTGTTCAGCAATCCAACACAATACCTCATTTTCTACACTCCCAACAA
ACTGTTCTCCATGCGAACCGGGCATGGAAGCGAGTCCCACGTGCCGCTGTGCGTATCCATTCA
TGGGAACACTCTACTTCCGGTCTCCTTCTTTCTCAGGGTTGTTCAACTCCACCAACTTCTCAATTC
TACAGAAGGCAATCGCGGATTTCTTTAAGAAGTTCAATTATCCTGTAGACTCCGTTGGTGTTCGGA
ACATAAGAGAGAATCCAACTGATCATCAGCTTCTAATAGATCTATTGGTCTTTCCATTGGGAAGAGA
GAGTTTTAATCAGACAGGAATGTCACTTGTTGGTTTTGCATTTAGCAACCAGACTTATAAGCCTCCT
CCGATATTTGGCCCTTATATTTTCAAAGCTGATTTGTACAAACAGTTCTCTGGGAGCGCTATGAAAA
TCGAAGAAGGTAAACTGGTAATCTGGATTAACGGCGATAAAGGCTATAACGGTCTCGCTGAAGTC
GGTAAGAAATTCGAGAAAGATACCGGAATTAAAGTCACCGTTGAGCATCCTGATAAACTGGAAGA
GAAATTCCCACAGGTTGCTGCAACTGGCGATGGCCCTGACATTATCTTCTGGGCACACGACCG
CTTTGGTGGCTACGCTCAATCTGGCCTGTTGGCTGAAATCACCCCTGACAAAGCATTCCAGGAC
AAGCTGTATCCATTTACCTGGGATGCCGTACGTTACAACGGCAAGCTGATTGCTTACCCTATCGC
TGTTGAAGCTTTATCACTGATTTATAACAAAGATCTGCTGCCTAACCCTCCAAAAACCTGGGAAGA
GATCCCTGCACTGGATAAAGAACTGAAAGCTAAAGGTAAGAGCGCACTGATGTTCAACCTGCAA

GAACCATACTTCACCTGGCCACTGATTGCTGCTGACGGGGGTTATGCATTCAAGTATGAAAACG
GCAAGTACGACATTAAAGACGTGGGCGTGGATAACGCTGGCGCTAAAGCTGGTCTGACCTTCC
TGGTTGACCTGATTAAAAACAAACACATGAATGCAGACACCGATTACTCCATCGCAGAAGCTGC
CTTTAATAAAGGCGAAACAGCTATGACCATCAACGGCCCATGGGCATGGTCCAACATCGACAC
CAGCAAAGTGAATTATGGTGTAACTGTACTGCCTACCTTCAAGGGTCAACCATCCAAACCATTCG
TTGGCGTGCTGAGCGCAGGTATTAACGCCGCCAGTCCTAACAAAGAGCTGGCAAAAGAGTTCC
TCGAAAACTATCTGCTGACTGATGAAGGTCTGGAAGCTGTTAATAAAGACAAACCACTGGGTGCC
GTAGCACTGAAGTCTTACGAGGAAGAGTTGGTGAAAGATCCACGTATTGCCGCCACTATGGAAA
ACGCCCAGAAAGGTGAAATCATGCCTAACATCCCTCAGATGTCCGCTTTCTGGTATGCCGTGCG
TACTGCGGTGATCAACGCCGCCAGCGGTCGTCAGACTGTCGATGAAGCCCTGAAAGACGCTC
AGACTTAG

Protein

MSSRTGASLLLILFFFQICSVSALTNGLDASALNALKSEWTTPPDGWEGSDPCGTNWVGITCQND
RVVSISLGNLDLEGKLPADISFLSELRILDLSYNPKLSGPLPPNIGNLGKLRNLILVGCSFSGQIPESIG
TLKELIYLSLNLNKFSGTIPPSIGLLSKLYWFDIADNQIEGELPVSNGTSAPGLDMLLQTKHFHFGKN
KLSGNIPKELFSSNMSLIHVLFDGNQFTGEIPETLSLVKTLTVLRLDRNKLIGDIPSYLNNLTNLNELYL
ANNRFTGTLPNLTSLTSLYTLDVSNNTLDFSPIPSWISSLPSLSTLRMEGIQLNGPIPISFFSPPQLQTV
ILKRNSIVESLDFGTDVSSQLEFVDLQYNEITDYKPSANKVLQVILANNPVCLEAGNGPSYCSAIQH
NTSFSTLPTNCSPCEPGMEASPTCRCAYPFMGTLYFRSPSFSGLFNSTNFSILQKAIADFFKKFNYPV
DSVGVRNIRENPTDHQLLIDLLVFPLGRESFNQTGMSLVGFAFSNQTYKPPPIFGPYIFKADLYKQFS
GSAMKIEEGKLVIWINGDKGYNGLAEVGKKFEKDTGIKVTVEHPDKLEEKFPQVAATGDGPDIIFWA
HDRFGGYAQSGLLAEITPDKAFQDKLYPFTWDAVRYNGKLIAYPIAVEALSLIYNKDLLPNPPKTWEE
IPALDKELKAKGKSALMFNLQEPYFTWPLIAADGGYAFKYENGKYDIKDVGVDNAGAKAGLTFLVDLI
KNKHMNADTDYSIAEAAFNKGETAMTINGPWAWSNIDTSKVNYGVTVLPTFKGQPSKPFVGVLSAG
INAASPNKELAKEFLENYLLTDEGLEAVNKDKPLGAVALKSYEEELVKDPRIAATMENAQKGEIMPNI
PQMSAFWYAVRTAVINAASGRQTVDEALKDAQT

**Nucleotide and amino acid sequences of EctoCARD1$^{HtriA}$-MBP**

EctoCARD1$^{HtriA}$-GSA linker-MBP

**Red letters denote His -> Ala mutation**

Nucleotide

ATGAGTTCAAGAACTGGAGCCTCTTTGCTCCTGATCTTGTTTTTCTTCCAAATTTGTTCTGTTTCGGC
ACTCACGAATGGTTTAGACGCTTCTGCTTTAAATGCTCTGAAGAGTGAATGGACCACTCCTCCTG
ATGGTTGGGAAGGCTCTGATCCTTGTGGAACCAATTGGGTTGGAATTACATGTCAAAATGACCGC
GTCGTTTCAATATCACTAGGTAACCTTGACTTGGAAGGAAAGCTTCCTGCCGATATATCGTTCTTGT
CCGAGTTGCGGATCTTGGATTTGTCATACAATCCTAAATTGTCCGGACCGCTTCCACCAAACATC
GGTAACCTTGGGAAGTTAAGGAACTTGATCCTTGTGGGATGTAGTTTCAGTGGTCAAATCCCTGA
GTCCATTGGAACTCTAAAAGAACTTATATATCTCTCCCTAAATTTAAATAAATTTAGTGGAACAATTCC
GCCTTCAATTGGCCTGTTATCAAAACTATATTGGTTTGATATAGCTGACAATCAGATTGAAGGAGAG
CTTCCAGTTTCTAATGGGACTTCTGCACCTGGACTTGACATGCTTCTTCAAACTAAG**GCT**TTT**GCT**
TTTGGAAAAAACAAGCTTTCAGGCAATATCCCAAAAGAACTTTTCAGCTCAAACATGAGTTTAATA
**GCT**GTCCTCTTCGATGGAAACCAATTCACGGGCGAAATTCCAGAGACCCTCAGTCTCGTTAAAA
CGTTGACGGTGTTACGCCTTGATAGGAACAAACTCATTGGAGATATTCCTTCATATCTTAACAATCT
CACAAATCTTAACGAACTGTACTTGGCCAACAACAGATTTACTGGTACTCTTCCAAATTTAACCAG
CTTGACCAGTCTCTACACATTGGATGTGAGCAATAACACTTTGGATTTCTCACCCATTCCATCATG
GATCTCTTCATTACCCTCCCTATCAACATTAAGGATGGAAGGGATCCAACTTAATGGTCCAATACC
AATCTCATTTTTCAGCCCTCCTCAGTTGCAGACTGTTATCTTAAAGCGAAATAGTATAGTGGAAAGC
TTAGACTTTGGTACAGACGTTAGCAGCCAGTTGGAGTTTGTGGATTTACAATACAATGAAATAACTG
ATTATAAACCATCAGCTAATAAAGTCCTCCAAGTAATATTGGCAAATAATCCAGTGTGTCTAGAGGC
GGGAAACGGGCCGAGTTACTGTTCAGCAATCCAACACAATACCTCATTTTCTACACTCCCAACA
AACTGTTCTCCATGCGAACCGGGCATGGAAGCGAGTCCCACGTGCCGCTGTGCGTATCCATTC
ATGGGAACACTCTACTTCCGGTCTCCTTCTTTCTCAGGGTTGTTCAACTCCACCAACTTCTCAATT
CTACAGAAGGCAATCGCGGATTTCTTTAAGAAGTTCAATTATCCTGTAGACTCCGTTGGTGTTCGG
AACATAAGAGAGAATCCAACTGATCATCAGCTTCTAATAGATCTATTGGTCTTTCCATTGGGAAGAG
AGAGTTTTAATCAGACAGGAATGTCACTTGTTGGTTTTGCATTTAGCAACCAGACTTATAAGCCTCC
TCCGATATTTGGCCCTTATATTTTCAAAGCTGATTTGTACAAACAGTTCTCTGGGAGCGCTATGAAA
ATCGAAGAAGGTAAACTGGTAATCTGGATTAACGGCGATAAAGGCTATAACGGTCTCGCTGAAGT
CGGTAAGAAATTCGAGAAAGATACCGGAATTAAAGTCACCGTTGAGCATCCTGATAAACTGGAAG
AGAAATTCCCACAGGTTGCTGCAACTGGCGATGGCCCTGACATTATCTTCTGGGCACACGACC
GCTTTGGTGGCTACGCTCAATCTGGCCTGTTGGCTGAAATCACCCCTGACAAAGCATTCCAGGA
CAAGCTGTATCCATTTACCTGGGATGCCGTACGTTACAACGGCAAGCTGATTGCTTACCCTATCG
CTGTTGAAGCTTTATCACTGATTTATAACAAAGATCTGCTGCCTAACCCTCCAAAAACCTGGGAAG
AGATCCCTGCACTGGATAAAGAACTGAAAGCTAAAGGTAAGAGCGCACTGATGTTCAACCTGCA

AGAACCATACTTCACCTGGCCACTGATTGCTGCTGACGGGGGTTATGCATTCAAGTATGAAAACG
GCAAGTACGACATTAAAGACGTGGGCGTGGATAACGCTGGCGCTAAAGCTGGTCTGACCTTCC
TGGTTGACCTGATTAAAAACAAACACATGAATGCAGACACCGATTACTCCATCGCAGAAGCTGC
CTTTAATAAAGGCGAAACAGCTATGACCATCAACGGCCCATGGGCATGGTCCAACATCGACAC
CAGCAAAGTGAATTATGGTGTAACTGTACTGCCTACCTTCAAGGGTCAACCATCCAAACCATTCG
TTGGCGTGCTGAGCGCAGGTATTAACGCCGCCAGTCCTAACAAAGAGCTGGCAAAAGAGTTCC
TCGAAAACTATCTGCTGACTGATGAAGGTCTGGAAGCTGTTAATAAAGACAAACCACTGGGTGCC
GTAGCACTGAAGTCTTACGAGGAAGAGTTGGTGAAAGATCCACGTATTGCCGCCACTATGGAAA
ACGCCCAGAAAGGTGAAATCATGCCTAACATCCCTCAGATGTCCGCTTTCTGGTATGCCGTGCG
TACTGCGGTGATCAACGCCGCCAGCGGTCGTCAGACTGTCGATGAAGCCCTGAAAGACGCTC
AGACTTAG

Protein

MSSRTGASLLLILFFFQICSVSALTNGLDASALNALKSEWTTPPDGWEGSDPCGTNWVGITCQND
RVVSISLGNLDLEGKLPADISFLSELRILDLSYNPKLSGPLPPNIGNLGKLRNLILVGCSFSGQIPESIG
TLKELIYLSLNLNKFSGTIPPSIGLLSKLYWFDIADNQIEGELPVSNGTSAPGLDMLLQTK**A**F**A**FGKNK
LSGNIPKELFSSNMSLI**A**VLFDGNQFTGEIPETLSLVKTLTVLRLDRNKLIGDIPSYLNNLTNLNELYLA
NNRFTGTLPNLTSLTSLYTLDVSNNTLDFSPIPSWISSLPSLSTLRMEGIQLNGPIPISFFSPPQLQTVI
LKRNSIVESLDFGTDVSSQLEFVDLQYNEITDYKPSANKVLQVILANNPVCLEAGNGPSYCSAIQHN
TSFSTLPTNCSPCEPGMEASPTCRCAYPFMGTLYFRSPSFSGLFNSTNFSILQKAIADFFKKFNYPVD
SVGVRNIRENPTDHQLLIDLLVFPLGRESFNQTGMSLVGFAFSNQTYKPPPIFGPYIFKADLYKQFSG
SAMKIEEGKLVIWINGDKGYNGLAEVGKKFEKDTGIKVTVEHPDKLEEKFPQVAATGDGPDIIFWAH
DRFGGYAQSGLLAEITPDKAFQDKLYPFTWDAVRYNGKLIAYPIAVEALSLIYNKDLLPNPPKTWEEI
PALDKELKAKGKSALMFNLQEPYFTWPLIAADGGYAFKYENGKYDIKDVGVDNAGAKAGLTFLVDLI
KNKHMNADTDYSIAEAAFNKGETAMTINGPWAWSNIDTSKVNYGVTVLPTFKGQPSKPFVGVLSAG
INAASPNKELAKEFLENYLLTDEGLEAVNKDKPLGAVALKSYEEELVKDPRIAATMENAQKGEIMPNI
PQMSAFWYAVRTAVINAASGRQTVDEALKDAQT
