## Supplemental Data 3 for "A copper-dependent, redox-based hydrogen peroxide perception in plants"

**Extended Data 3. List of CARD1 homologs identified from the NCBI database.**

>AT5G49760 CARD1 [Arabidopsis thaliana]
MSSRTGASLLLILFFFQICSVSALTNGLDASALNALKSEWTTPPDGWEGSDPCGTNWVGITCQNDRVVSI
SLGNLDLEGKLPADISFLSELRILDLSYNPKLSGPLPPNIGNLGKLRNLILVGCSFSGQIPESIGTLKEL
IYLSLNLNKFSGTIPPSIGLLSKLYWFDIADNQIEGELPVSNGTSAPGLDMLLQTKHFHFGKNKLSGNIP
KELFSSNMSLIHVLFDGNQFTGEIPETLSLVKTLTVLRLDRNKLIGDIPSYLNNLTNLNELYLANNRFTG
TLPNLTSLTSLYTLDVSNNTLDFSPIPSWISSLPSLSTLRMEGIQLNGPIPISFFSPPQLQTVILKRNSI
VESLDFGTDVSSQLEFVDLQYNEITDYKPSANKVLQVILANNPVCLEAGNGPSYCSAIQHNTSFSTLPTN
CSPCEPGMEASPTCRCAYPFMGTLYFRSPSFSGLFNSTNFSILQKAIADFFKKFNYPVDSVGVRNIRENP
TDHQLLIDLLVFPLGRESFNQTGMSLVGFAFSNQTYKPPPIFGPYIFKADLYKQFSDVEVSSKSSNKSIL
IGAVVGVVVLLLLLTIAGIYALRQKKRAERATGQNNPFAKWDTSKSSIDAPQLMGAKAFTFEELKKCTDN
FSEANDVGGGGYGKVYRGILPNGQLIAIKRAQQGSLQGGLEFKTEIELLSRVHHKNVVRLLGFCFDRNEQ
MLVYEYISNGSLKDSLSGKSGIRLDWTRRLKIALGSGKGLAYLHELADPPIIHRDIKSNNILLDENLTAK
VADFGLSKLVGDPEKTHVTTQVKGTMGYLDPEYYMTNQLTEKSDVYGFGVVLLELLTGRSPIERGKYVVR
EVKTKMNKSRSLYDLQELLDTTIIASSGNLKGFEKYVDLALRCVEEEGVNRPSMGEVVKEIENIMQLAGL
NPNSDSATSSRTYEDAIKGSGDPYGSESFQYSGNFPASKLEPQ
>XP_002323702.2 leucine-rich repeat receptor protein kinase HPCA1
[Populus trichocarpa]
MGLGGGTARLLFLLTFFTSGIHLIFSDTDPSDAAALQSLKKQWQNTPPSWGQSHDPCGAPWEGVTCSNSR
ITALGLSTMNLKGKLSGDIGGLTELRSLDLSFNTNLTGSLTPRFGDLLKLNILILAGCGFSGSIPDELGN
LAELSFLALNSNNFSGGIPPSLGKLSKLYWLDLADNQLTGPIPISKNTTPGLDLLLNAKHFHFNKNQLSG
SIPPELFSSDMVLIHVLFDGNQLEGNIPSTLGLVQTLEVLRLDRNALSGKVPKNLNNLSSLNELNLAHNK
LIGPLPNLTKMDALNYVDLSNNSFYSSEAPDWFSTLPSLTTLVIEHGSLHGTLPSKVFSFPQIQQVLLRN
NALNGSFNMGDSISTQLQLVDLQNNQISSVTLTADYTNTLILVGNPVCTALSDTNYCQLQQQSTKPYSTS
LANCGSKMCPPEQKLSPQSCECAYPYEGTLYFRAPSFRELSNVNMFHSLEMSLWGKLGLTPGSVFLQNPF
FNVDDYLQVQVALFPPTDKYFNRSEIQSIGFDLTNQTYKPPKDFGPYYFIASPYPFPDASRGSSMSTGVV
VGIGIGCGLLVMSLVGVGIYAIRQKKRAEKAIGLSKPFASWAPSGKDSGGVPQLKGARWFSYEELKRCTY
NFTESNEIGSGGYGKVYRGMLSDGQVVAIKRAQQGSMQGGLEFKTEIELLSRVHHKNLVGLVGFCFEQGE
QMLVYEYMPNGTLRECLSGKSGIYLDWRRRLRIALGSARGLAYLHELANPPIIHRDVKSTNILLDENLTA
KVADFGLSKLVSDSSKGHVSTQVKGTLGYLDPEYYMTQQLTEKSDVYSFGVVMLELIAAKQPIEKGKYIV
REVRMAMDRNDEEHYGLKEIMDPGLRNMGGNLVGFGRFLEVAMQCVEESATERPTMSEVVKAIEMILQND
GVNTNSTTSASSSATDFGASRGGGPLRHPYNHDVVAAKNKVDVVDNINNNNAFDYSGGYTLSAKVEPK
>XP_012446614.1 leucine-rich repeat receptor protein kinase HPCA1
[Gossypium raimondii]

MGSAVWLFLVVVLFHIYIVAAETDPDDNSSLRAVMSEWKNVPSNWGRGDPCDDKWVGIGCTGSRVTSLNL
PNMKLEGRLVGDIFFLSELKELDLSYNKGLRGILPPAVQNLKKLENLILVGCGFSGQIPDTIGSLPQLRI
LSLNSNAFSGNIPPSIGNLSTLNWLDMADNQLEGEIPVSNGSTTPGLDWLIHTKHFHFGLNKLSGPIPRK
LFSSDMTLIHVLFENNMLSGPLPLTLGLVKTLEVVRFDNNSLEGDLPLNLNNLTRVQDLYLSNNKLTGPL
PNLTGMSSLNTLYLSNNSFDSSDVPSWFPTLLSLTTLMMESTQLKGQIPASFFNLPQLQTVVLKQNELDG
SFDIGPSFSNQLQIINLQGNSITSFNNTGGPISFDIVLVDNPVCQETGAGANDYCSLPQPDSSSVYTTAP
MNCVPNSCGSGQISSPRCICAYPYTGTLQFRGLYFSNLRNGTPYESLEQNLTQFFRFPELLVDTVSLSNP
RMDQHLYLLLDLYLFPYGQDRFNTSGISKIASAFSSQDYKPPEQYFGPYVFTGAEYEYFSDGPAHSNKSS
AGIAIGAAVGASVLFILLVVAGIYAYRQRKRADRATKESNPFAHWDPKKSSGSIPQLKGARCFSFEELKK
YTKKFSEANDIGSGGYGKVYRGTLPTGELVAIKRAQQGSMQGGLEFKTEIELLSRVHHKNVVSLLGFCFE
RGEQMLVYEYIPNGSLSDSLSGKSGIRLDWPRRLKIALGAAKGVAYLHELANPPIIHRDIKSTNILLDER
LNAKVADFGLSKPMGDSEKGHVSTQVKGTMGYLDPEYYMTQQLTEKSDVYSFGVLMLEIITARRPIERGK
YIVREMRMSMDKTKSLYNLQQILDPAIGFGTSSKGLERFVELAMRCVEESGADRPTMGEVVKEIENIMQM
DGMNPNAESASSSATYEDATKGADLHPYDNESFAYSGAFPHSAAKIEPH

>XP_006470792.2 leucine-rich repeat receptor protein kinase HPCA1
[Citrus sinensis]

MGVKRKVFLLSVYLQFLIIAAVTNDNDFVILKALKDDIWENEPPNWKNNDPCGDNWEGIGCTNSRVTSIT
LSGMGLKGQLSGDITGLTELHTLDLSNNKDLRGPLPTTIGNLKKLSNLMLVGCSFSGPIPDSIGSLQELV
LLSLNSNGFSGRVPPSIGNLSNLYWLDLTDNKLEGEIPVSDGNSPGLDMLVRAKHFHFGKNQLSGSIPEK
LFRPDMVLIHVLFDSNNLTGELPDTLGLVKSLEVVRFDRNSLSGPVPSNLNNLTSVNELYLSNNKLTGAM
PNLTGLSVLSYLDMSNNSFDASEVPSWFSSMQSLTTLMMENTNLEGQIPANLFSIPHLQTVVMKTNELNG
TLDLGTSYSENLLVNLQNNRISAYTERGGAPAVKLTLIDNPICQELGTAKGYCQLSQPISPYSTKQKNCL
PAPCNANQSSSPNCQCAYPYTGTLVFRSLSFSDLGNTTYYEILEQNVTTSFQSTYKLPIDSISLSYPHKN
NFEYLELSIQFFPSGQERFNRTGVSSVGFVLSNQIYSPPPLFGPMFFNGDQYQYFAESGGSNKSTSIGVI
IGAAAAGCVVLLLLLFAGVYAYHQKRRAEKANEQNPFAHWDMNKSSGSIPQLKGARCFSFEEVKKYTNNF
SDANDVGSGGYGKVYKGTLPNGQLIAIKRAQQGSMQGGQEFKMEIELLSRVHHKNLVSLLGFCFDRGEQM
LIYEFVPNGSLGDSLSGKNGIRLDWIRRLKIALGAARGLSYLHELANPPIIHRDIKSSNILLDERLNAKV
ADFGLSKSMSDSEKDHITTQVKGTMGYLDPEYYMTQQLTEKSDVYSFGVLMLELLTGRRPIERGKYIVRE
IRTVMDKKKELYNLYELIDPTIGLSTTLKGFEKYVDLALKCVQESGDDRPTMSEVVKDIENILQQAGLNP
NAESASSSASYEDASKGNFHHPYCNEEGFDYGYSGGFPTSKIEPQ

>XP_014617332.1 leucine-rich repeat receptor protein kinase HPCA1
isoform X1 [Glycine max]

MGERVIVLILLFFTHLLIVFTKTSPQDSAALLALVNEWQNTPPNWDGTDPCGAGWDGIECTNSRITSISL
ASMDLSGQLTSDIGSLSELLILDLSYNKKLTGPLPNDIGNLRKLRNLLVINCGFTGPIPVTIGNLERLVF
LSLNSNGFTGPIPAAIGNLSNIYWLDLAENQLEGPIPISNGTTPGLDMMHHTKHFHFGKNKLSGNIPSQL
FSPEMSLIHVLFESNRFTGSIPSTLGLVKTLEVVRFDDNVLSGPVPLNINNLTSVRELFLSNNRLSGSPP

NLTGMNSLSYLDMSNNSFDQSDFPPWLPTLPALTTIMMENTKLQGRIPVSLFSLQQLQTVVLKNNQLNGT
LDIGTSISNNLDLLDLQINFIEDFDPQIDVSKVEIILVNNPICQETGVPQTYCSITKSNDSYSTPPDNCV
PVPCSLDQTLSPECKCAYPYEGTLVLRAPSFSDLENKTIFVTLESSLMESFQLHKKPVDSISLSNPRKNI
YQYLELTLKIFPLGQDRFNRTGISDIGFLLSNQTYKPPPMFGPYYFIADEYENYVDNSEGPVTSNRKSSN
TGIIAGAGGGGAALLVLVLLACVYAISQKKKTKKSTGNNNPFEQWDPHDSNSSIPQLKGARRFSFEEIQN
CTKNFSQVNNIGSGGYGKVYRGTLPNGQLIAVKRAQKESMQGGLEFKTEIELLSRVHHKNLVSLVGFCFD
QGEQMLIYEYVANGTLKDTLSGKSGIRLDWIRRLKIALGAARGLDYLHELANPPIIHRDIKSTNILLDER
LIAKVSDFGLSKPLGEGAKGYITTQVKGTMGYLDPEYYMTQQLTEKSDVYSFGVLLLELITARRPIERGK
YIVKVVKGAIDKTKGFYGLEEILDPTIDLGTALSGFEKFVDIAMQCVEESSFDRPTMNYVVKEIENMLQL
AGSSPIFSASASVSTSSSYNNATKISLHPYNNEYFDSSVVLPRA

>XP_003594434.2 leucine-rich repeat receptor protein kinase HPCA1 isoform X1 [Medicago truncatula]

MGERTLVFLLFLFSYLLVVVVTKTSNDDYLALSTLKYEWKNVPPSWEDSEDPCGDHWEGIECSNSRVITI
SLSSMDLSGQLSSEIGSLSELQILVLSYNKDLTGPLPAEIGNLKKLTNLQLINCGFTGPIPDTIGNLQRL
VFLSLNSNRFSGRIPPSIGNLSNINWLDLAENQLEGPIPVSNGTTPGLDMLHKTKHFHFGKNKLSGNIPP
QLFSSDMSLIHVLFESNQFTGTIPSTLGFVQKLEVVRLDNNILSGPLPININNLTNVRELLVSKNRLSGP
LPDLTGMNVLSYLDVSNNSFDRSDFPLWLSTLQSLKTIMMEDTQLQGPIPVSLFSLVQLHTVMLKNNNLN
GTLDIGTAISDQLGVLNLQTNFIEDFDPQIDVSKVEIILVNNPVCQETGVKRTYCSIAKNNDTYTTPLNN
CVPVECNKNQILSPKCKCAYPYTGTLTLRAPSFSDVRNKTVFAMLEFTLMESFRLHEKPVDSVSLSNPRK
NAYQYLDLSLEIFPSGQDSFNRTGISGIGFMLSNQTYKPPAETFGPYYFIADKYEHYLNDSVIEGPVKSS
KSSHIGIIAGAAAGGCVLVLLLLLAVVYGFRQKNKAKRAAKKSNLFEQWGPDESNSSIPQLKGARRFTFE
EIQNYTKKFAEASYVGSGGYGKVYRGALLNGQLIAVKRAQKESIQGGLEFKTEIELLSRVHHKNLVSLIG
FCFEQGEQILVYEYVVNGTLTDALSGKSGIRLDWIRRLKIALGASRGLDYLHEHANPPIIHRDVKSTNIL
LDERLNAKVSDFGLSKPLGDGAKGYITTQVKGTMGYLDPEYYMTQQLTEKSDVYSFGVLMLELITARRPI
ERGKYIVKVIKNAMDKTKELYGLKEIIDPVIDFKASLSSFEKFIDLAMKCVEDSSSSRPSMNYAFKEIEN
MLMLTGTNPNAESAPSSSSYNESGNSMHPYENEYFDSSVILPRA

>XP_019079160.1 leucine-rich repeat receptor protein kinase HPCA1 [Vitis vinifera]

MDSRLILVSLLIVFIQISATWARTNTDDATALVALKDLWENYPPSWVGFDPCGSSWEGIGCYNQRVISII
LTSMGLKGGLSGDLDQLSELQILDLSYNKNLTGNIPASIGSLKKLTNLILVGCSFSGPIPDTIGSLTELV
FLSLNSNSFSGGIPPSIGNLSKLYWLDLADNQLTGTIPISNGSTPGLDKLTHTKHFHFGKNRLSGSIPPK
LFSSNMILIHLLLESNRLTGSIPSTLGLLKTLEVVRLDGNSLSGPVPSNLNNLTEVKDLFLSNNKLTGTV
PDLTGMNSLNYMDMSNNSFDVSNVPSWLSTLQSLTTLTMENTNLKGAIPASLFSLPQLQTVSLRNNIING
TLDFGAGYSSQLQLVDLQKNYIVAFTERAGHDVEIILVENPICLEGPKNEKYCMTSQPDFSYSTPPNNCV
PSVCSSDQIPSPNCICAYPYMGTLVFRAPSFSNLGNSSYYISLEQRLMQSFQSQQLPVDSVFLADLMKDS
NNYLQVSLKVFPHGRDRFNRTGISMVGFALSNQTFKPPSTFGPFYFNGEQYQYFEEVSLSLEPNKSSNTG

IIIGAAVGGSLLVLLLLFAGVYAFRQKRRAERATEQSNPFANWDESKGSGGIPQLKGARRFTFEEIKKCT
NNFSDVNDVGSGGYGKVYRATLPTGQMVAIKRAKQESMQGGLEFKTEIELLSRVHHKNVVSLIGFCFQLG
EQILIYEYVPNGSLKESLSGRSGIRLDWRRRLKVALGSARGLAYLHELADPPIIHRDIKSNNILLDEHLN
AKVGDFGLCKLLADSEKGHVTTQVKGTMGYMDPEYYMSQQLTEKSDVYSFGVLMLELISARKPIERGKYI
VKEVKIAMDKTKDLYNLQGLLDPTLGTTLGGFNKFVDLALRCVEESGADRPTMGEVVKEIENIMQLAGLN
PITESSSASASYEESSTGTSSHPYGSNSAFDSSAGYPPSTVEPK

>XP_004230391.1 probable leucine-rich repeat receptor-like protein kinase At5g49770 [Solanum lycopersicum]

MVVSQAMIPRILLCFLFVLIHVLSIAARTNPDDSAALQSLKDSWQNVPPNWVGADPCGSSWDGIGCRNSR
VVSITLSSMSLEGQLSGDIQGLAELETLDLSYNKELKGSLPQSIGKLTKLSNLILVGCGFSGPIPDTIGS
LTRLVFLSLNSNNFIGGIPATVGYLTELYWLDLADNKLTGTIPVSNGSSPGLDLLVHTKHFHFGKNQLSG
EIPAGLFHSNLSLIHLLVENNKFTGNIPDTLGLVQTMEVLRLDRNSLSGSVPQNLNNLTHVNELHMSNNN
FNGLLPNLTGMNVLNYLDMSNNSFNASDFPSWIPNLISLTSLVMENTGLQGTVPASLFSLYQLQTVILRN
NKLNGSLTIDTTYSNQLQLIDVQRNLIESFTQRPGYPFQIMLAGNPFCNEGGDGTQDYCVKTQQTETYST
PPENCLPTDCSSNRVSSPTCKCAFPYTGNIVFRAPSFSNLGNRTTYETLQKSLMQTFQNRQLPVESVSLS
NPTKNLDDYLVIHLQVFPSTQDFFNRTGVSGIGFVLSNQTFKPPSSFGPFFFIGEGYKYFDGASSESKNS
SSTGIIIGAAVGGSVIAIIALIIGVYAFRQKKRAEDAAKRSDPFASWDSNKHSGAVPQLTGARFFSFEEL
KKWTNNFSETNDIGCGGYGKVYRGTLPNGELVAIKRALQGSMQGAHEFKTEIELLSRVHHKNVVGLAGFC
FDQAEQMLVYEYIPNGTLKDGLSGKTGIRLDWMRRLRIAVGAARGLQYLHDLVNPPIIHRDIKSNNILLD
DRLNARVADFGLSKLLGDSDRGHITTQVKGTMGYMDPEYYMTNQLTEKSDVYSFGVVLLEIVTGKVPIEK
GRYIVREVKTAMDRSKDMYNLQDILDPAVRAGATPRSLEKFVDLALKCVEEEGANRPSMSEVVKEIENIM
EMAGLNPNADSASSSATYEGPNKGMNHPYTDESLFVYSGAYPNSKVEPK

>XP_015619793.1 probable leucine-rich repeat receptor-like protein kinase At5g49770 [Oryza sativa Japonica Group]

MGVSPWIIIFLLIVLVQAFVASADTNAQDTSGLNGLAGSWGSAPSNWAGNDPCGDKWIGIICTGNRVTSI
RLSSFGLSGTLSGDIQSLSELQYLDLSYNKNLNGPLPSTIGTLSKLQNLILVGCGFTGEIPKEIGQLSNL
IFLSLNSNKFTGSIPPSLGGLSKLYWFDLADNQLTGGLPISNATSPGLDNLTSTKHFHFGINQLSGSIPS
QIFNSNMKLIHLLLDNNKFSGSIPSTLGLLNTLEVLRFDNNAQLTGPVPTNLKNLTKLAEFHLANSNLTG
PLPDLTGMSSLSFVDMSNNSFSASDAPSWITTLPSSLTSLYLENLRISGEVPQSLFSLPSIQTLRLRGNR
LNGTLNIADFSSQLQLVDLRDNFITALTVGTQYKKTLMLSGNPYCNQVNDDVHCKATGQSNPALPPYKTT
SNCPALPPTCLSTQQLSPTCICSVPYRGTLFFRSPGFSDLGNSSYFIQLEGTMKAKFLNLSLPVDSIAIH
DPFVDTNNNLEMSLEVYPSGKDQFSEQDISGIGFILSNQTYKPPSNFGPYYFLGQTYSFANGALQTSKSN
TNHIPLIVGASVGGAAVIAALLALTICIARRKRSPKQTEDRSQSYVSWDIKSTSTSTAPQVRGARMFSFD
ELKKVTNNFSEANDIGTGGYGKVYRGTLPTGQLVAVKRSQQGSLQGNLEFRTEIELLSRVHHKNVVSLVG
FCFDQGEQMLVYEYVPNGTLKESLTGKSGVRLDWKRRLRVVLGAAKGIAYLHELADPPIIHRDIKSSNVL
LDERLNAKVSDFGLSKLLGEDGRGQITTQVKGTMGYLDPEYYMTQQLTDRSDVYSFGVLLLEVITARKPL

ERGRYVVREVKEAVDRRKDMYGLHELLDPALGASSALAGLEPYVDLALRCVEESGADRPSMGEAVAEIER
IAKVAGAGGAAAAESAASDSMSYAASRTPRHPYGGGGGDSASEYSGGGLPSMRVEPK
>XP_021302278.1 probable leucine-rich repeat receptor-like protein kinase At5g49770 isoform X1 [Sorghum bicolor]
MELSPWLVFSFGFLAQALVILADTNVQDTAGLNGIKDSWNKKPSNWVGTDPCGDKWIGIDCTGDRVTSIR
LSSLGLSGSLSGDIQSLSELQTLDFSYNKDLGGPLPASIGSLSNLENLIIVGCSFSGEIPKELGQLSKLI
FLSMNSNKFSGSIPPSLGRLSKLYWFDLADNKLSGELPVFDGTNPGLDNLTNTKHFHFGINQLSGTIPSQ
IFNSHMKLIHLLLDNNNFTGSIPSTLGLLNTLEVLRFDNNYQLTGSVPSNINNLTKLAELHLENNKLNGP
LPDLTGMIALSFVDMSNNSFNASDVPSWFTTLPSLTSLYLENLRVTGQLPQDLFSLPAIQTLRLRGNRFN
GTLTIGSDFSTQLQLIDLRDNDISQITVGGSQYNKQLILVGNPICSSGSNEKYCTPPGQSNQATPPPYST
AKNCSGLPPPCLSGSGQLLSPSCACAVPYRGTLFFRSPSFSDLSNGSYWGQLESGIKAKYLSLSLPVDSV
AIHDPSVNSVNNLQVALEVFPGGKTMFSEQDISDIAFVLSNQTYKPPSVFGPYYFNGQQYSFANELLIPS
KSKSNNLPLIIGVSAGGAVLVAGVVALVICVARRKKKKRPKQNEERSQSFVSWDMKSTSGGSSSIPQLRG
ARMFSFDELRKITNNFSEANDIGNGGYGKVYRGTLPTGQLVAVKRSQQGSLQGSLEFRTEIELLSRVHHK
NVVSLVGFCLDQAEQILVYEYVPNGTLKESLTGKSGVRLDWRRRLRVVLGAAKGVAYLHELADPPIVHRD
IKSSNVLLDERLNAKVSDFGLSKPLGDDGRGQVTTQVKGTMGYLDPEYYMTQQLTEKSDVYSFGVLMLEV
ATARKPLERGRYIVREMKAALDRTKDLYGLHDLLDPVLCAAPSAPEGMEQYVDLALRCVEEAGADRPSMG
EVVSEIERVLKMAGGAGPESASNSMSYASRTPRHPYGGDSPFADYSSAGLPSARVEPK
>Brara.J00581.1.p pacid=30613296 transcript=Brara.J00581.1 locus=Brara.J00581 ID=Brara.J00581.1.v1.3 annot-version=v1.3
MSSRIGAFMLLILLCFQFLSVSALTNGFDASALQALKADWTKYPENWVGADPCGTNWVGITCTNDRVVSI
SLGNLDVEGKLSSDIASLTELQILDLSYNTELTGPLPSNIGQLKKLKNLILVACSFSGQIPESIGDLEQL
IYLSLNLNQFSGRIPASIGRLSNLYWFDIADNQIEGTIPVSNGTSSPGLDMLLETKHFHFGKNKLSGVIP
ETLFSSKMTLIHVLFDGNNFTGDIPDTLSLVKTLTVLRLDRNKLTGNIPTSLNNLTNLQELYLANNEFTG
SLPNLTSLTSLYTLDVSNNTLEFSPIPSWISSLRSLATLRMEGINLNGSIPNSFFSPPQLQTVILKRNRI
NSALDFGTSYSNQLEFVDLQYNDIDVYTQPSSNTRIQVILANNPVCQEQGNSPSYCSAIPHNTSYSTIPT
TCSPCDHGREASPSCRCAHPFTGTFNFRAPSFSGLFNSTNFEILQKDITGFFNKFSYPVDSVAVRNIREN
TTDHQLLIDLLVFPLGRESFNETGMLLVNFAFSNQTYKPPPIFGPYIFIADPYTQFSDGGGFKSSNMGVI
IGAAVGCAVLLLLLLTLAGVYALCQRKRADRATDQNNPFAKWSTSKSSIDAPQLMGAKSFTFEELKKCTDN
FSEANDVGGGGYGKVYRGILPSGQLIAIKRAQQGSLQGGLEFKTEIELLSRVHHKNVVRLLGFCFDRSEQ
MLVYEYIPNGSLRDSLSGKSGIRLDWIRRLRIALGSGKGLAYLHELADPPIIHRDIKSNNILLDENLTAK
VADFGLSKLVGDPEKTHVTTQVKGTMGYLDPEYYMTNQLTEKSDVYGFGVVMLELLTGKSPIEKGKYVVR
EVKMKMNKSRSLYDLQELLDTTIIASSSNLKGFDKYVDLALRCVEEEGVNRPSMGDVVKEIENIMQLAGL
NPNGDSASTSATYEDAIKGSGDPYGKDSFQYSGNFPASKLEPQ
>QJR84046.1 CARD1-like protein 1 [Phtheirospermum japonicum]

MCWRIYLSLLLVLVHVFGISALTYTDDFVALKSLKDVWNNVPPNWDGADPCGSHWDGIGCNANNRVVTIT
LASINISGQLSSDIEGLSELQTLDLSYNKGMTGALPTAIGNLRQLSSLILVGCGFSGPIPPSVGSLQQLV
YLSLNSNNFVGEIPATIGNLSKLYWLDLADNKLSGPIPVSRGTTPGLDMLVTTKHFHFGNNQLSGEIPSQ
LFSSNLTLIHLLLENNQLTGRIPSSLALVQTLEIVRLDRNLLNGSVPVNLNSLTSVNELYLANNGLTGPL
PNLTGMGLLHYVDLSNNSFDPTYVPAWFSSLSSLTSLIMENTLIEGQLPVSLFSLSQLQIVTLKNNRING
TLNMGSSYSSQLQRIDLQNNFVDAFTERAGFNVSVILVGNPICDEGVTQSYCTIPPQSNTTYTTPTENCT
PSPCSSEQVSSPTCQCSYPYSGMLYFRAPSFYNYGNATVFESLQNKMMSTFTSHQQPVDSVSVSNPTKNI
DNYLLLSLQVFPSGEDHFNRTGISRIGFFLSNQTFKPPEGFGPFYFIANNYPYFEGLKEGSKRSSNIGVI
IGAAAGGSILFLLLLIAGIYAIRQKKRAETATKKSDPFALWDTNSNSGGVPQLRGAKNFSFDEIKKCTNN
FSEMNEIGSGGYGTVHRGTLRNGQLVAIKRSQKGSLQGGVEFKNEIELLSRVHHKNVVSLVGFCFDQGEH
MLVYEYITNGSLKDSLNGKTGIRLDWTRRLRIALGAARGVQYLHELANPPIIHRDIKSNNILLDERLNAK
VADFGLSKPMGEPEKGHITTQVKGTMGYLDPEYYMTQELTEKSDVYSFGVLLFELLTSRAPIVKGKYVVR
EVKETMDKTKNLYNLESILDPIVASNMAPGSVEKFVDLALRCVQELGVNRPTMSEVVKEIENIMEMAGLN
PHNESASTSGSYEGAKKELSHPYTNESLLSHSGAYSPSI

>QNJ34483.1 CARD1-like protein 1 [Striga asiatica]

MSWGISFFLLVALLQVFGIAALTNTEDFVALKSLKDVWENVPPNWDGADPCGSGNGWDGINCDANNRVVT
ITLVSFNISGQLSSDIKGLSELQILDLSYNRGLTGALPSAIGNLKKLTSLILVGCGFSGPIPPSIGSMQE
LVYLSLNLNNFVGEIPATIGNLPNLYWLDLADNRLSGTIPVSTRTTPGLDMLFKTNHFHFGNNRLSGEIP
SQLFNSNMTLIHLLLEGNQLTGRIPSSLALVQSLEVIRLDRNSLSGSVPNNLNSLTTVQELSLANNRLTG
PFPNLTGMNLLHSVDLSNNSFDATDVPSWLSSSTSLTTLIMDNTGIHGQLPVSLFSLPQLQTVELKNNRI
NGTLNIGPSPSSQLQRIDLQNNFIDGYTERAGYNASVQIVLVGNPICDEGATQSYCRIPPQTNNTTTYQT
PPENCTPTSCNSDRIQSPTCRCAYPYSGALFFRAPSFSSYGNATIFESLRQKLMSTFQTHRQPVDSVSLS
NPTRNMENYLQLTLQIFPAGQDHFNRTGVSRIGFIMSNQTFKPPPGFGPFYFNADTYPYFAGPNEGSNRP
SNIHVIIGAAVGASILLLLLLIAGVYAFRQKRRAETAAKKSDPFASWHSNSNSGAVPQLRGARSFSFEEI
KKCTDNFSDNNEIGSGGYGKVYRGTLPNGQLVAIKRTKPGSTQGGVEFKNEIELLSRVHHKNVVCLVGFC
FDHGEFMLVYEYITNGSLKDSLTGKTGIRLDWTRRLRIAIGAARGIQYLHELANPPIIHRDIKSNNILLD
DRLNAKVADFGLSKPMGEPEKGHITTQVKGTMGYLDPEYYMTQELTEKSDVYSFGVLLFELLTSRSPIVN
GKYIVREVKELMDKTKNLYNLEPVLDPIVASNMAPGSVEKFVDLALRCVQELGVSRPTMNEVVKEIENIM
ELSGLNPHNESASTSSSYEGKMKEHGHPYTYESLSSHNEVFSPSI

>XP_004137665.1 probable leucine-rich repeat receptor-like protein kinase At5g49770 isoform X1 [Cucumis sativus]

MSPVETLLLFAFFYAGIDTAGSFTDPRDSAALESLRNEWQNTPPSWGASIDPCGTPWEGVACINSRVTAL
RLSTMGLKGKLGGDIGGLTELKSLDLSFNKDLTGSISPALGDLQNLSILILAGCGFSGSIPEQLGNLSNL
SFLALNSNNFTGTIPPSLGKLSNLYWLDLADNQLTGSLPVSTSETPGLDLLLKAKHFHFNKNQLSGSISP
KLFRSEMVLIHILFDGNKFSGNIPPTLGLVKTLEVLRLDRNSLAGTVPSNLNNLTNINELNLANNKLTGP
LPNLTQMSSLNYVDLSNNSFDSSEAPEWFSNLQSLTTLIIEFGSMRGSVPQGVFSLPQIQQVKLKKNAFS

DTFDMGDKVSEQLQLVDLQNNNISHFTLGSRYTKTLMLIGNPVCSTDVTLSNTNYCQVQDQPVKPYSTSL
ASCLSKSCSPDEKLSPQSCECTYPFEGTLYFRAPSFRDLSNVTLFHSLEFSLWKKLDLTPGSVSIQNPFF
NVDDYLQMQLALFPSDGKYFNRSEIQRIGFYLSNQTYKPPHEFGPFYFIASPYGFADTTKGTSISPGVII
GVAIGCAFLVLGLIGVGIYAIWQKKRAEKAIGLSRPFASWAPSGNDSGGAPQLKGARWFSYDELKKCTNN
FSMSNEVGSGGYGKVYRGMLVDGQAVAIKRAQQGSMQGGLEFKTEIELLSRVHHKNLLGLVGFCFEQGEQ
MLVYEFMPNGTLRDSLSGKSGINLDWKRRLRIALGSARGLAYLHELANPPIIHRDVKSTNILLDEHLNAK
VADFGLSKLVSDNEKGHVSTQVKGTLGYLDPEYYMTQQLTEKSDVYSFGVVMLELLTGKLPIEKGKYVVR
EVRMLMNKSEEEYYGLKQIMDVTILNNTTTIIGLGRFLELAMRCVEESAGDRPTMSEMVKAIESILQNDG
INTNTTSASSSATDFGASRNAPRHPYNDPIPKKDAHDSNSFDYSGGYTLSTKVEPK

>XP_009391655.2 leucine-rich repeat receptor protein kinase HPCA1 isoform X1 [Musa acuminata AAA Group]

MGTLVFLLFVFLASLQTSSGSTDAQDAAALLSLMTQWQNTPPSWGKTDDPCGTPWEGVRCSNSRVTSLTL
STMGIKGTLGDDIGQLSELKILDLSYNTMLGGTLTPNIGNLMELTILILVGCSFNGNIPDELGSLGNLSY
LALNSNQFTGSIPASLGKLSNLNWFDIADNQLTGPLPISTETSPGLDQLVRTQHFHFNKNQLSGPIPEKL
FSSDMTLLHVLFDGNNFTGKIPDSIGLVQKIQVLRLDRNALSGPVPSNINNLTHVKELNLANNRLTGMMP
NLTGMNSLNYVDLSNNTFDASETPAWFSELQSLRALVIESGGLYGEVPKELFSFTQLQQVILDNNEFNGT
LDMGNSISQQLQIVNFKNNKLTGVAHDASYDRTLILIGNPLCDWLSMTKFCSLRQEPAIPSYSTSLAECA
ANLCPPDQSLNPQNCKCVYPYEGVMFFRAPLFRDVTNSTLFQSLESSLWKKLDLPPGSVFLQNPFFNNDS
YLQVQVKFFPSSGMYFNRSEILQIGFKLSNQIYKPPEIFGPYYFKAFQYPFPGVEGKSPIAIGLIIGIAV
GCALLVIGLLLITIYALRQRKQAQRAIKLSKPFASWAHSGDEIVDAPQLKGARWFSYDELRRCTDNFSVS
NEIGSGGYGKVYKGMLPGGQVVAIKRAQQGSMQGGHEFKTEIELLSRVHHKNLVALVGFCFDEGEQMLVY
EFIPNGTLREGLSGKSGILLDWRRRLRIALGSARGLAYLHELADPPIIHRDVKSSNILLDEKLNAKVADF
GLSKLVSDNEKGHISTQVKGTMGYLDPEYYLTQQLSDKSDVYSFGVVMLELMTAKQPLEKGKYIVREVKM
AIDADDEEFYGLKELMDHAIQNAAYLIAFRKFAELALRCLEESAGDRPSMSDVVKEIEIMLNADGLSTNS
NSASSSATDFGYAKGVPKHPYDSHSRKDVSSSNSFEYSGGYTFSTKPEPK

>NP_001344074.1 Probable leucine-rich repeat receptor-like protein kinase At5g49770 precursor [Zea mays]

MNSPKMELSPWLVFSLGFLAQALVILADTNAQDASGLNGIQDSWNKKPSNWRVGTDPCGDKWNGISCTTS
RVTAIRLSSLGLSGSLSGDIQSLSELQTLDFSYNKDLGGPLPASIGSLSNLENLILVGCSFSGEIPKELG
QLTKLRFLSLNSNKFSGSIPASLGRLSNLYWFDLADNKLSGGLPVFDGTNPGLDNLTNTKHFHFGVNQLS
GTIPSQIFNSNMTLIHVLLDNNNFTGRIPPTLGLLNTLEVMRFDNNYQLTGPVPSNINNLTKLAELHLEN
NQLTGPLPDLTGMIALSFVDMSNNSFNASGVPSWFTTLPSLTSLYLENLRVTGQLPQALFSLPAVQTLRL
RGNRFNGTLTIGSDYSTQLQLIDLRDNQISQITVGGSQYNKQLILVGNPICSPGTGSSEKYCASPGQSNQ
AAPPPYSTPMNCSGLPPPCLSDQLVSPGCVCAVPYRGTLFFRSPSFSDLSNGSYWGQLETGIRAKFRSLS
VPVDSVALHDPSVNSVNNLQLALEVFPSGKTQFSEQDISDIGFILSNQTYKPPSVFGPYYFLGQPYSFAN
VVLIPSKSKANNRLPLIVGASVGGAVLVAIVLALVTIVARRKKRPKQNEERSQSFVSWDMKSTSGSSVPQ

LRGARTFNFDELRKITSNFSEANDIGNGGYGKVYRGTLPSGQLVAVKRCQQGSLQGSLEFRTEIELLSRV
HHKNVVSLVGFCLDQAEQILVYEYVPNGTLKESLTGKSGVRLDWRRRLRVLLGAAKGIAYLHELADPPIV
HRDIKSSNVLLDERLNAKVSDFGLSKPLGEDGRGQVTTQVKGTMGYLDPEYYMTQQLTDKSDVYSFGVLM
LEMATARKPLERGRYIVREMKVALDRTKDLYGLHDLLDPVLGSSPSALAGLEQYVDLALRCVEEAGADRP
SMGEVVGEIERVLKMAGGPGPESASNSMSYASRTPRHPYGGDSPFDHSNSGLPSARVEPK

>XP_021592398.1 leucine-rich repeat receptor protein kinase HPCA1 isoform X1 [Manihot esculenta]

MQNFKLKKNGEFLRFYPEMNPKNLVFLLVASLQIWSIAALTNNADLTVLKAVMDMWENPPPSWEGTDPCG
DQWDGIKCINSRVTSITLSSMGLKGQLSGDITNLPELLILDLSYNKDLRGPLPASIGNLKKLRNLILLGC
SFSGPIPSSIGSLQQLLFLSLNSNGFSGPIPPSIGNLSELYWLDLADNKLDGSIPVSTGTTPGLDMLVKT
KHFHLGKNQLSGEIPPKLFSSDMTLLHVLFDDNKLTGSIPSTLGLVQTLEVIRFDRNSLTGPVPSNLNNL
TSVSELFLSNNGLTGPLPNLTGMSFLSYLDMSNNSFDASDFPPWTSTLQSLTTLILEGTQLQGQIPSSFF
SLANLQNVVLSNNRLNGTLDIGTVNSGQLQLIDLQSNFISDYTPQPGQNQVYVILVNNPVCQETGVKASF
CTDLRPNSSYVTLPNNCVPVPCGSNKISSPNCNCAYPYTGVLVFRAPSFSDLGNINVYVSLQKDLMDSFK
SNQLPVDSVSLSNPRKDSSEYLDLNLQVFPSEKDNFNRTVISEIGFLLSNQTFKPPDFFGPYYFIADAYQ
YFAGEATGSNNSSNTGIIIGAVVGGSALVLLLLLAGLYAYRQKKRAERATELNNPFANWDSTKSNGAGVP
QLKGARLFSFEELRKYTNNFSEANDIGSGGYGKVYRGTLPNGELIAIKRAQQESMQGGLEFKTEIELLSR
VHHKNLVSLLGFCFDRGEQMLVYEFVPNGSLSDSLSAGKSGIRLDWVRRLKIALGAARGLVYLHELANPP
IIHRDIKTNNILLDERLNAKVADFGLSKPMSDTEKGHITTQVKGTLGYLDPEYYMTQQLTEKSDVYSFGV
VMLELLTGRKPIERGKYIVREVRMAMDRTKDLYNLHELLDPGIGLETTLKGLDKFVDLAMECVKESGADR
PMMGDVVKEIETILQLAGLNPNAESASTSASYEEAGKGSTHPYNKESFYYSGAFPPSKLEPK
